# Permanent lymphocyte subset elimination upon a single dose of AAV-delivered depletion antibody dissects immune control of chronic viral infection

**DOI:** 10.1101/2024.06.19.599723

**Authors:** Anna Lena Kastner, Anna-Friederike Marx, Mirela Dimitrova, Tiago Abreu-Mota, Yusuf I. Ertuna, Weldy V. Bonilla, Karsten Stauffer, Ingrid Wagner, Mario Kreutzfeldt, Doron Merkler, Daniel D. Pinschewer

## Abstract

To interrogate the role of specific immune cells in infection, cancer and autoimmunity, immunologists commonly use monoclonal depletion antibodies (depletion-mAbs) or genetically engineered mouse models (GEMMs). To generate a tool that combines specific advantages and avoids select drawbacks of the two methods we engineer adeno-associated viral vectors expressing depletion-mAbs (depletion-AAVs). Single-dose depletion-AAV administration permanently eliminates lymphocyte subsets in mice while avoiding accessory deficiencies of GEMMs such as marginal zone defects in B cell-deficient animals. Depletion-AAVs can be used irrespective of the animals’ genetic background, and multiple depletion-AAVs can readily be combined. Exploiting depletion-AAV technology, we show that B cells are required for unimpaired CD4 and CD8 T cell responses to chronic viral infection. Importantly, CD8 T cells fail to suppress viremia when B cells are depleted, and they only help resolving chronic infection if antibodies suppress viral loads. Our study positions depletion-AAVs as a versatile tool for immunological research.

## Introduction

At repeated occasions over the past decades technological advancements have redefined the way biomedical research is conducted. In immunology, for example, the advent of monoclonal antibodies (mAbs) has marked a new era^1^. Besides their widespread use as detection reagents, one key application of mAbs in basic research consists in the selective depletion of individual hematopoietic cell types in experimental animals^2–6^. Accordingly, mAb-based depletion is widely used to this day to investigate the contribution of individual lymphocyte subsets in a broad range of biological processes. One main limitation of this approach consists, however, in its intrinsically transient nature. Antibody half-life and the replenishment of depleted populations from hematopoietic precursors determine the durability of the experimental intervention. Hence an effective long-term depletion of cell types with an intrinsically high turnover rate can require that mAbs be re-administered at short interval throughout the experiment^7–10^. This is both expensive and inconvenient for research personnel. Moreover, repeated administrations have the potential to compromise animal welfare and under certain conditions such as in high-containment facilities can be impossible to implement.

In this regard, genetically engineered mouse models (GEMMs) offer key advantages. Introduced into basic biomedical research several decades ago^11–15^, CRISPR/Cas-mediated genome engineering has recently revolutionized the versatility of this approach, allowing for a more efficient, faster and cheaper generation of GEMMs^16^. However, also GEMMs have clear limitations and shortcomings. The choice of parental mouse strain for GEMM generation can decide over the resulting phenotype^17^, and transfer of a modified genetic locus into another genetic background can require years of backcrossing. Accordingly, it has been proposed that outbred mouse colonies of defined genetic diversity may represent a better surrogate of the human population^18–20^. Such models are, however, inherently incompatible with GEMM technology. The same applies to feral mice, which may represent a better surrogate of the adult human immune system than animals from specific pathogen-free housing conditions^21,22^. Additional frequent shortcomings of GEMMs consists in compensatory mechanisms but also in accessory deficiencies that emerge when specific lymphocyte subsets are lacking throughout ontogeny. Examples consist in the MHC class I-restricted cytotoxicity found in CD8-deficient but not CD8 T cell-depleted mice^23^, and in the virtual absence of a marginal zone in B cell-deficient mice^24,25^. Such accessory deficiencies are particularly problematic when investigating mechanisms of protective immune defense against infection. The exquisite susceptibility of B cell-deficient mice to vesicular stomatitis virus (VSV) induced disease, for example, was long thought to be due to the absence of a virus-neutralizing antibody (nAb) response^26^, but may actually relate primarily to the role of B cells in establishing protective macrophage compartments^27,28^.

Approximately 300 million people are affected by persistent viral diseases such as HIV and hepatitis B and C virus (HBV, HCV), with wide-ranging medical and socio-economical consequences^29–31^. Unlike for HCV, a functional cure for HIV or HBV carriage remains unavailable, emphasizing the need to better understand how the immune system can prevail against persisting viral intruders. Infection of mice with lymphocytic choriomeningitis virus (LCMV) represents the prototype small animal model for such investigations. Accordingly, LCMV research in mice has contributed several fundamental concepts^32^ including T cell exhaustion^33–35^, which has subsequently been validated in the aforementioned human diseases as well as in cancer^36–38^. While LCMV research has taken center stage in establishing the key role of CD8 T cells in resolving primary viral infection^39–43^ shortcomings of GEMMs as outlined above have long compromised investigations into the role of B cells in resolving chronic infection^25,44–48^. Recent evidence obtained from GEMMs with more targeted alterations in B cells support a critical role for B cells themselves in resolving chronic LCMV infection^25,49–51^. The relative contribution of CD8 T cells and B cells to viral load control and importantly also the interdependence of their antiviral efficacy in the chronic infection context remains, however, ill-defined.

Here we develop adeno-associated viral vectors (AAVs)^52–54^ delivering depletion antibodies (depletion-AAVs) as a novel approach for the permanent depletion of lymphocyte subsets in mice. Depletion-AAVs combine the versatility of classical antibody depletion strategies with the permanent nature of GEMMs. They can readily be used in mice of diverse genetic background, allow for the combined depletion of more than one lymphocyte subset and enable an unperturbed ontogenic development of secondary lymphoid organs. Accordingly, the exploitation of depletion-AAV technology has offered new insights into the synergism of CD8 T cells and B cells in resolving chronic viral infection.

## Results

### A single-dose of AAV-vectored antibody affords long-term lymphocyte subset depletion

AAVs allow for long-term expression of their encoded transgenes including antibodies^53,54^. Thus, we hypothesized that depletion-AAVs would allow for the permanent elimination of lymphocyte subsets in mice without a need for vector re-administration (Fig. 1A). As a proof-of-concept we engineered an AAV vector encoding the CD4 depletion antibody YTS191 (AAV-αCD4; Fig. S1A-B). This rat antibody was converted to a mouse IgG2a format (YTS191-mIgG2a; referred to as αCD4), offering optimal antibody-dependent cellular cytotoxicity (ADCC) and reducing the immunogenicity of the AAV-vectored antibody^8,55^ while allowing its discrimination from endogenous IgG2b and IgG2c in C57BL/6 mice^56^. We administered to mice intramuscularly (i.m.) either a single dose of AAV-αCD4 or of an AAV-vector encoding an antibody of irrelevant specificity (AAV-ctrl). A third group of mice was given a single dose of αCD4 mAb intraperitoneally (i.p.). Subsequently, the animals were infected with lymphocytic choriomeningitis virus (LCMV) to study immune responses (Fig. 1B,G). Recipients of AAV-αCD4 exhibited serum αCD4 concentrations that on average remained at levels >500 µg/ml and the animals were devoid of circulating CD4 T cells for an observation period of >100 days (Fig. 1C,D, Fig. S2B). In contrast αCD4 mAb-treated mice showed gradual recovery of a circulating CD4 T cell compartment from around day 21 after depletion onwards, as expected^57^. Analysis of the spleen on day 102 confirmed quasi-complete depletion of CD4 T cells in AAV-αCD4-treated mice but a largely restored CD4 T cell compartment in αCD4 mAb-treated controls (Fig. 1E-F). In keeping with virtually complete depletion of CD4 T cells from spleen and lymph nodes (Fig. S2C-D) AAV-αCD4-treated mice did not mount a germinal center (GC) B cell response to LCMV infection and failed to produce LCMV-specific IgG serum antibodies, which are T help-dependent (Fig. 1H-I)^58^.

**Figure 1:**
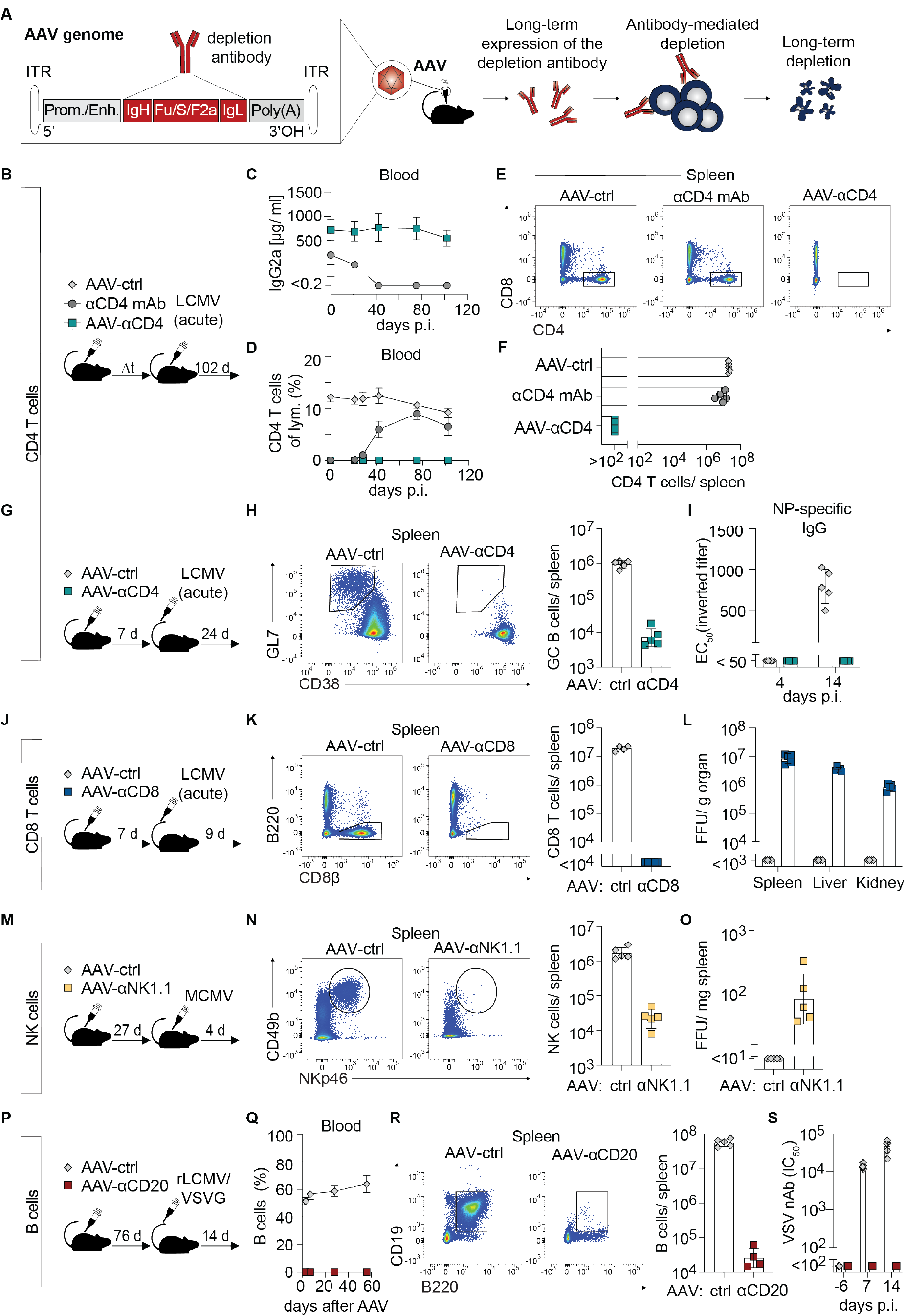
A single-dose of AAV-vectored antibody affords long-term lymphocyte subset depletion. A: Schematic of the strategy for AAV-mediated lymphocyte subset depletion. B-F: We administered AAV-αCD4 or AAV-ctrl to mice one week prior to acute LCMV infection (B). A separate group of mice was treated with anti-CD4 depletion antibody (αCD4 mAb) as recombinant protein on d-3 and d-1 of infection. The concentration of the depletion antibody in serum (C) and the percentage of CD4 T cells amongst peripheral blood lymphocytes (D) were monitored over time. Representative FACS plots (E; pre-gated on Ter119^−^CD19^−^ live lymphocytes) and absolute numbers of CD4 T cells (F) on d102 p.i. in spleen. Gating strategies for all data sets are summarized in Fig. S2A. G-I: Mice were treated with AAV-αCD4 or AAV-ctrl, followed by acute LCMV infection one week later (G). Representative FACS plots and absolute numbers of GC B cells (GL7^+^CD38^−^ CD19^+^CD8^−^CD45.2^+^ live lymphocytes) in spleen 24 days p.i. (H). LCMV NP-specific serum IgG titers at the indicated time points after infection (I). J-L: We treated mice with AAV-αCD8 or AAV-ctrl, followed by acute LCMV infection one week later (J). Representative FACS plots (left) and absolute CD8 T cell counts (gated as CD8β^+^B220^−^ live lymphocytes, right) in spleen (K), and viral loads in spleen, liver and kidney on d9 after LCMV infection (L). M-O: We treated mice with AAV-αNK1.1 or AAV-ctrl, followed by MCMV infection 27 days later (M). Representative FACS plots and absolute numbers of NK cells (NKp46^+^CD49b^+^CD3^−^ live lymphocytes) (N) and viral loads (O) in spleen on d31 after AAV-treatment i.e. d4 after MCMV infection. P-S: We treated mice with AAV-αCD20 or AAV-ctrl, followed by rLCMV/VSVG infection 76 days later (P). The frequency of B cells (B220^+^CD19^+^) amongst peripheral blood lymphocytes was monitored (Q). Representative FACS plots (left) and absolute numbers of B cells (B220^+^CD19^+^ live lymphocytes; right) in spleen on day 90 after AAV treatment (R). VSV-nAb titers were determined at the indicated time points after rLCMV/VSVG infection (S). Symbols in (C,D,Q) represent the mean±SD of five mice per group. In panels (F,H,I,K,L,N,O,R,S) symbols represent individual mice and bars indicate the mean±SD. For each data set one representative of two independent experiments is shown.

To further assess the versatility of AAV-vectored long-term leukocyte subset depletion we developed depletion-AAVs expressing either CD8 T cell-, B cell-, NK cell-or neutrophil-depleting antibodies, all of them in a mouse IgG2a format (Fig. 1J-S, S1B, S2E-J). The CD8 T cell-depleting AAV encoded the CD8α-specific antibody YTS169 (AAV-αCD8; Fig. S1B). We validated the AAV-αCD8 in the context of acute LCMV infection (Fig. 1J), the clearance of which is strictly CD8 T cell-dependent^39–43^. Single-dose AAV-αCD8 treatment reduced the splenic CD8 T cell compartment to below detection limits (Fig. 1K) and prevented viral clearance from spleen, liver and kidney (Fig. 1L), whereas AAV-ctrl-treated mice were virus-free by day 9 after infection. To assess long-term NK cell depletion, we treated mice with a single dose of AAV-αNK1.1 expressing the widely used mAb PK136 (AAV-αNK1.1; Fig. S1B). For at least one month after administration the CD49b^+^NKp46^+^ NK cell compartments in spleen, lymph nodes and bone marrow were virtually abolished (Fig. 1M-N, S2E-F). Accordingly, single-dose AAV-αNK1.1 treatment impaired the NK cell-dependent control^59^ of mouse cytomegalovirus infection (MCMV), when virus challenge was performed four weeks after administration of AAV-αNK1.1 (Fig. 1O).

To deplete B cells, we vectorized the CD20-targeting antibody 18B12 (AAV-αCD20; Fig. S1B). B cells were undetectable in blood of AAV-αCD20 treated mice for at least 60 days (Fig. 1P-Q), and their numbers in spleen as well as in lymph nodes on day 90 were over a 1000-fold reduced (Fig. 1R and Fig. S2G). Mature IgD^+^ B cells in bone marrow were also depleted (Fig. S2H). To test the impact of long-term AAV-αCD20-mediated B cell depletion on antibody responses, mice were immunized with a recombinant LCMV (rLCMV/VSVG) expressing the glycoprotein (GP) of vesicular stomatitis virus (VSVG) instead of its own GP^60,61^. This vector elicits a potent T-independent neutralizing antibody (nAb) response but unlike its parent VSV does not cause disease in immunocompromised mice^62^. A single dose of AAV-αCD20 abolished VSV-nAb responses to rLCMV/VSVG immunization conducted 76 days later, whereas AAV-ctrl-treated mice mounted nAb responses that were >100-fold over technical background (Fig. 1S). An AAV delivering the Ly-6G/Ly-6C-specific mAb RB6.8C5 (AAV-αGr1; Fig. S1B) was also developed and resulted in the complete depletion of neutrophils over an observation period of more than two weeks (Fig. S2I-J), demonstrating that also granulocyte subsets are susceptible to depletion-AAVs. Taken together, these studies demonstrated that depletion-AAVs represent a versatile tool for the targeted long-term depletion of a broad range of leukocyte populations.

### AAV-delivered decoy receptor blocks IL-33 signaling to antiviral CD8 T cells in mice of different genetic backgrounds

Next, we tested whether AAV vector-based delivery could also be exploited for long-term blockade of cytokine signals in immune responses. We designed an AAV vector expressing a soluble decoy receptor (AAV-ST2-Fc^63^; Fig. S1C-D) for interleukin-33 (IL-33), a cytokine that is sensed by activated CD8 T cells through their receptor ST2. IL-33 represents a key driver of potent effector CD8 T cell responses to a wide range of viruses including LCMV^64^. To test whether IL-33 signaling could be blocked, we administered AAV-ST2-Fc to WT mice and two weeks later infected them with LCMV to assess antiviral CD8 T cell responses (Fig. 2A). As controls we used AAV-ctrl-treated WT mice as well as two GEMMs. *T1-Fc^tg^* mice express an ST2-Fc decoy receptor as a transgene and *IL-33^−/−^* mice lack a functional IL-33 gene. AAV-ST2-Fc-treated mice exhibited a 5-fold and 3-fold lower CD8 T cell response to the immunodominant H-D^b^-restricted epitopes GP33 and NP396 than AAV-ctrl treated animals, respectively (Fig. 2B-E). Moreover, KLRG1^+^CD127^−^ GP33- and NP396-specific effector CD8 T cells were 9-fold reduced. Importantly, these responses in AAV-ST2-Fc-treated mice were indistinguishable from those of *T1-Fc^tg^* and *IL-33^−/−^* mice, attesting to efficient AAV-ST2-Fc-mediated blockade of IL-33 signaling. Subsequently, we determined whether AAV-ST2-Fc can serve as a tool to assess the importance of IL-33 – ST2 signaling in antiviral CD8 T cell responses of BALB/c mice (Fig. 2F). AAV-ST2-Fc treatment resulted in a 3-fold lower expansion of CD8 T cells specific for the immunodominant H-2L^d^-restricted NP118 epitope and in a 5-fold reduction of the corresponding effector CD8 T cell response (Fig. 2G-H), closely mimicking effects of IL-33 deficiency on NP-specific CD8 T cell responses in C57BL/6 mice. These results indicated that AAV-vectored delivery can be exploited to block cytokine signaling pathways, yielding results comparable to cytokine knock-out and decoy receptor-transgenic models, and that AAV-based cytokine blockade can readily be used in mice of different genetic backgrounds.

**Figure 2:**
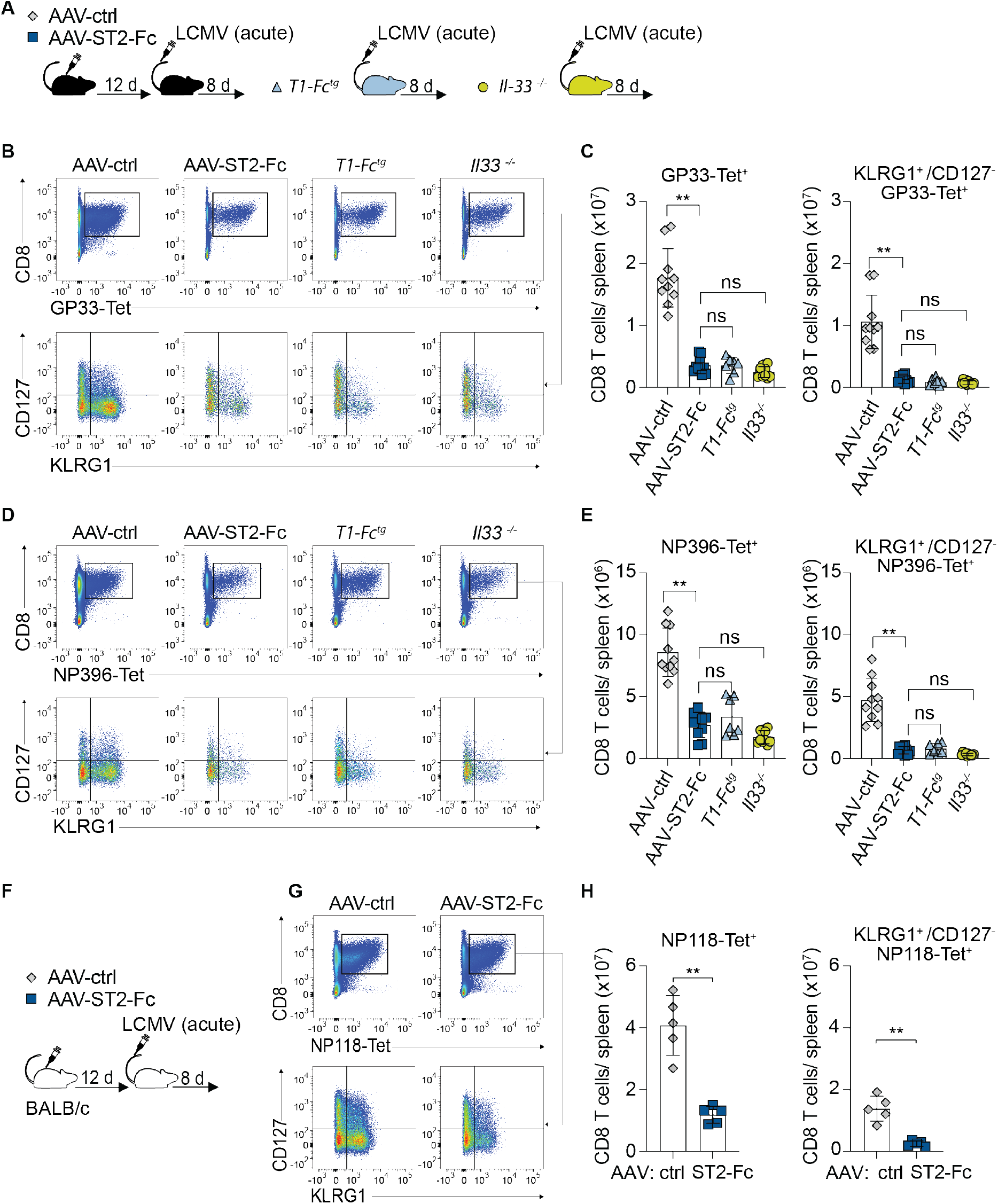
AAV-delivered decoy receptor blocks IL-33 signaling to antiviral CD8 T cells in mice of different genetic backgrounds. A-E : We administered an AAV-ctrl or an AAV-ST2-Fc to C57BL/6 WT mice (A). Two weeks later we infected these animals as well as groups of *T1-Fc^tg^* and *IL-33*^−/−^ control mice with LCMV (acute infection). Representative FACS plots of GP33-Tet^+^ (B) and NP396-Tet^+^ (D) CD8 T cells (top rows, pre-gated on CD4^−^B220^−^ live lymphocytes) and their phenotype (bottom rows, pre-gated on Tet^+^CD8^+^CD4^−^B220^−^ live lymphocytes) on d8 p.i. in spleen. Absolute numbers of GP33-Tet^+^ (C) and NP396-Tet^+^ (E) CD8 T cells (left) and of the KLRG1^+^CD127^−^ effector subset contained therein (right). F-H: We treated BALB/c mice with either AAV-ctrl or AAV-ST2-Fc two weeks prior to LCMV infection (acute setting; F). Representative FACS plots of NP118-Tet^+^ CD8 T cells (top row, pre-gated on CD4^−^B220^−^ live lymphocytes) and their phenotype (bottom row, pre-gated on NP118-Tet^+^CD8^+^CD4^−^B220^−^ live lymphocytes) in spleen 8 days p.i. (G). Absolute numbers of NP118-Tet^+^ CD8 T cells (left) and NP118-Tet^+^KLRG1^+^CD127^−^ effector CD8 T cells (right; H). Symbols in (C,E,H) represent individual mice and bars indicate the mean ± SD. (C,E) show combined data from two independent experiments, analyzed by one-way ANOVA followed by Dunnett’s post-test. Data in (H) are representative of two independent experiments analyzed by unpaired t test.**: p <0.01.

### B cell elimination by depletion-AAV maintains splenic marginal zone

B cells are required for a normal development of secondary lymphoid organs and by consequence, B cell-deficient mice exhibit defects in their splenic microarchitecture including a paucity in marginal zone macrophages^24,25^. We hypothesized that a depletion-AAV targeting B cells, which is administered to adult animals after completion of splenic development, may leave the splenic microarchitecture intact. Upon administration of AAV-αCD20 to mice we determined the extent of B cell depletion and the integrity of the splenic marginal zone over time. Untreated WT mice and B cell-deficient *J_H_T* mice served as controls (Fig. 3A). More than 95 % of the splenic B cell compartment was depleted within four days after AAV-αCD20 administration, reaching >98 % by d7 and quasi-complete depletion (>99 %) at later time points (Fig. 3B-D). Analogous B cell depletion kinetics were observed in lymph nodes (Fig. S3A), and in bone marrow the mature (IgD^+^) B cell compartment underwent >99 % depletion within four days after AAV-αCD20 administration (Fig. S3B). Interestingly, splenic marginal zone B cells (CD21^hi^CD23^lo^) exhibited somewhat delayed depletion kinetics as compared to transitional and follicular B cells (Fig. 3E). To determine whether AAV-αCD20 affected the integrity of the splenic microarchitecture, we detected on histological sections SIGN-R1^+^ marginal zone macrophages (MZM), CD169^+^ metallophilic marginal zone macrophages (MMM) and F4/80**^+^** red pulp macrophages alongside with CD19^+^ B cells (Fig. 3F). As expected^25^, B cell-deficient *J_H_T* GEMM mice exhibited profound marginal zone defects evident in a virtual absence of MMMs and MZMs. In remarkable contrast and despite virtually complete ablation of the B cell compartment, mice depleted of B cells by an administration of AAV-αCD20 mice 45 days earlier displayed an intact marginal zone as judged by the density and distribution of MMMs as well as MZMs, which resembled untreated WT mice. In summary, AAV-αCD20 treatment ablated the B cell compartment by >99% but unlike in B cell-deficient GEMMs left the splenic microarchitecture intact.

**Figure 3:**
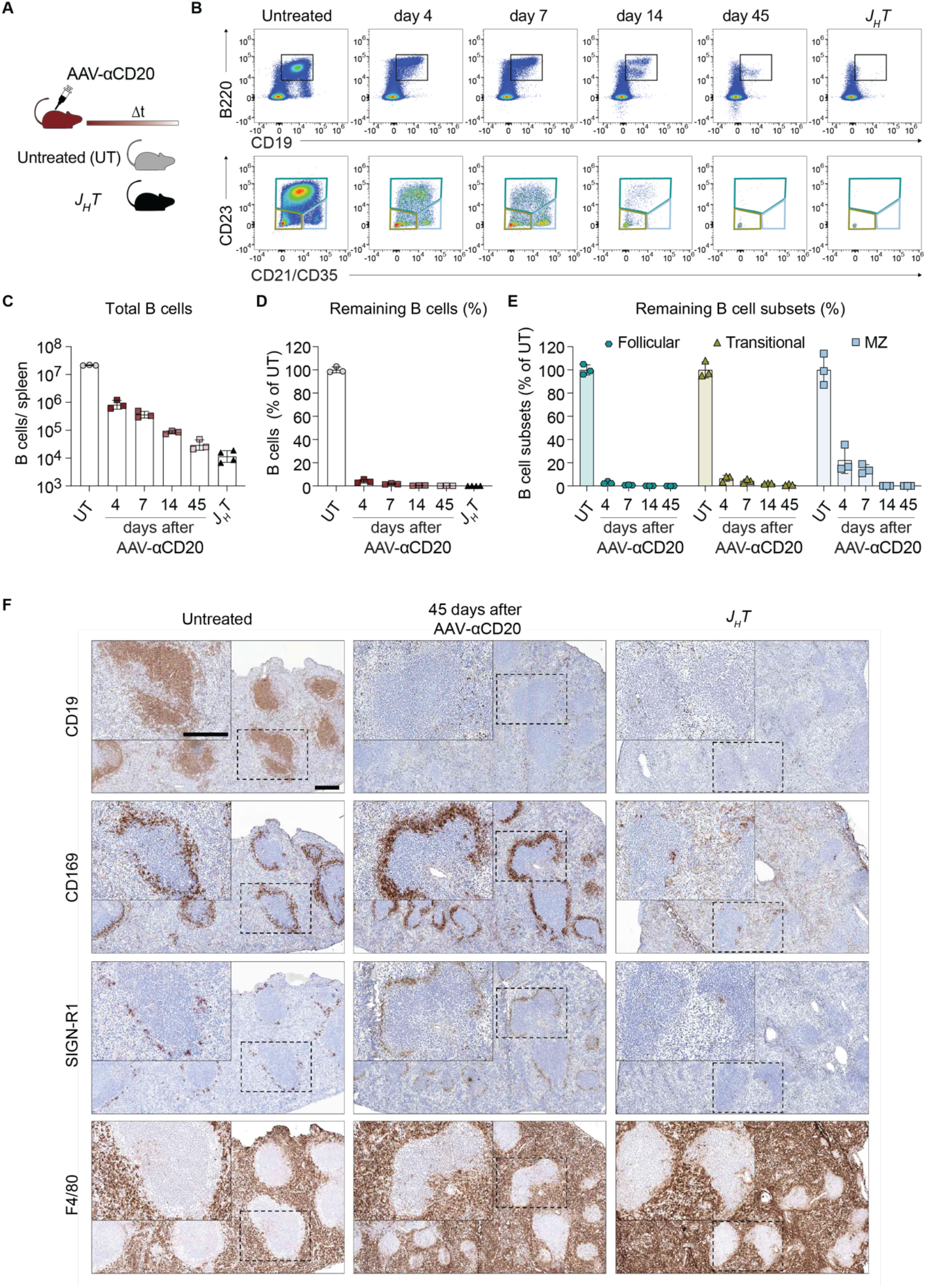
B cell elimination by depletion-AAV maintains splenic marginal zone. A-F : We administered AAV-αCD20 to mice and monitored the depletion of B cells and the integrity of the splenic microarchitecture over time (A). Untreated WT and *J_H_T* mice are shown for comparison. Representative FACS plots of B cells (top row, pre-gated on live lymphocytes) as well as of marginal zone (MZ; CD21/CD35^+^CD23^−^), transitional (CD21/CD35^−^CD23^−^) and follicular B cells (CD21/CD35^+^CD23^+^; bottom row, all pre-gated on B220^+^CD19^+^ live lymphocytes) in spleen (B) and their quantification (C). Depletion efficiency of B cells (D) and of individual B cell subsets (E) in spleen was calculated using untreated control mice as reference. Representative histological spleen sections (F) were stained for CD19 (B cells), CD169 (MOMA-1; metallophilic marginal zone macrophages), SIGN-R1 (ERTR9; marginal zone macrophages) and F4/80 (red pulp macrophages). Magnification bars: 200 µm. Symbols in (C,D,E) represent individual mice and bars indicate the mean±SD. One representative data set from two independent experiments is shown.

### Depletion-AAVs are effective during chronic LCMV infection

Chronic LCMV infection reduces the efficiency of antibody-mediated lymphocyte subset depletion, which has been linked to suppression of Fcγ receptor functions^65,66^. More complete depletion can, nevertheless, be achieved by increasing the dose of depletion antibody^67,68^. Chronic LCMV infection is also associated with hypergammaglobulinemia^69^, which can shorten serum antibody half-life^70,71^ and may further contribute to inefficient antibody-mediated depletion. To address this possibility, we administered an IgG2a mAb of irrelevant specificity to mice, which were either 19 days into chronic LCMV infection or uninfected (Fig. 4A). Monitoring the mAb’s concentration in serum we calculated a half-life of 7.5 days in uninfected mice, consistent with published results^72,73^, whereas in chronically LCMV-infected animals the antibody’s half-life was only 1.5 days i.e. ∼5-fold reduced (Fig. 4B). We hypothesized that the drastically reduced half-life of passively administered mAb may contribute to the relative inefficiency of antibody-mediated lymphocyte subset depletion in chronically LCMV-infected mice^65,66,68^ and that a continuous production of high amounts of depletion antibody by depletion-AAV technology might overcome these limitations. We established chronic LCMV in mice followed by AAV-αCD20, AAV-αCD8 or AAV-αCD4 treatment two weeks later (Fig. 4C). Within one week after depletion-AAV administration the blood of mice was virtually devoid of B cells, CD8 T cells or CD4 T cells, respectively (Fig. 4D and S4A). Analysis of the spleen two weeks after depletion-AAV administration corroborated virtually complete ablation of CD4 and CD8 T cells in the respective treatment groups (Fig. 4E-F). Total splenic B cell numbers in AAV-αCD20-treated animals were ∼17-fold below AAV-ctrl-treated mice (Fig. 4E-F). Non-class-switched B cells were ∼100-fold reduced, which was in keeping with virtually complete B cell depletion in uninfected mice. In remarkable contrast, class-switched (IgM^−^IgD^−^) and GC (GL7^+^CD38^−^) B cells, which overlapped to the largest extent (Fig. S4B), were only ∼4-fold and ∼3-fold reduced, respectively (Fig. 4G-H).

**Figure 4:**
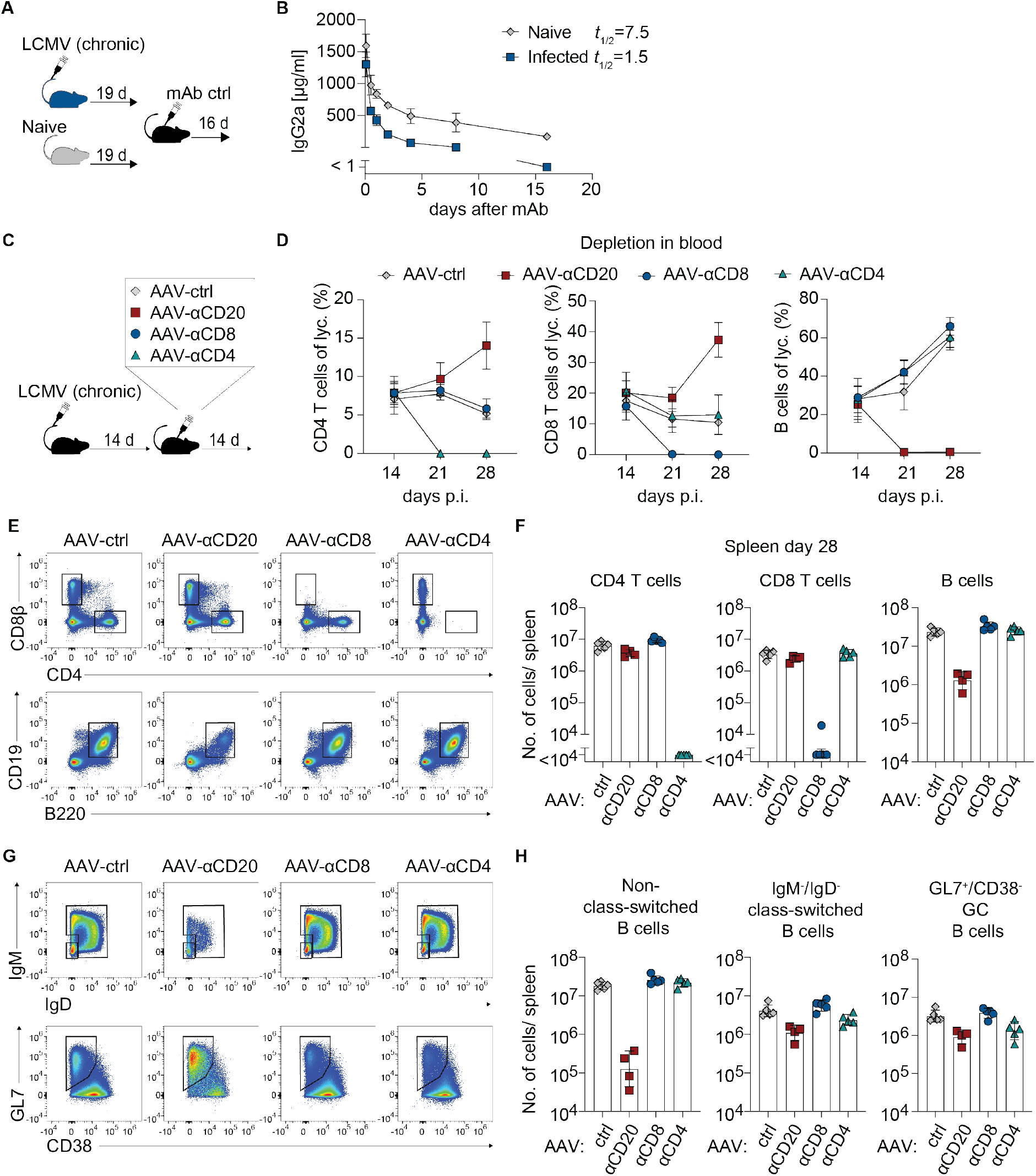
Depletion-AAVs are effective during chronic LCMV infection. A-B: We infected mice with LCMV (chronic setting) or left them uninfected, and19 days later administered 2 mg IgG2a mAb of irrelevant specificity (A). We monitored IgG2a concentration in serum over time (B) and calculated its half-life (*t*_1/2_) based on a one-phase decay model. C-H: We infected mice with LCMV (chronic setting) and treated them with AAV-ctrl, AAV-αCD20, AAV-αCD8 or AAV-αCD4 two weeks later (C). Frequencies of CD4 T cells, CD8 T cells and B cells amongst peripheral blood lymphocytes were monitored over time (D). Representative FACS plots of CD4 T cells and CD8 T cells (top row, pre-gated on B220^−^ live lymphocytes; E) and of B cells (bottom row, pre-gated on live lymphocytes; E) as well as their quantification (F) on d28 p.i. in spleen. Representative FACS plots of non-class-switched (either IgM^+^, IgD^+^ or IgM^+^IgD^+^) and class-switched (IgM^−^IgD^−^) B cells (top row) and GC (GL7^+^CD38^−^) B cells (bottom row, both pre-gated on B220^+^CD19^+^ live lymphocytes) in spleen on d28 p.i. (G) and their quantification (H). Symbols in (B) represent the mean±SD of three mice per group. Symbols in (D) represent the mean±SD of four (AAV-αCD20) to five mice per group (all other groups). In panels (F,H) symbols represent individual mice and bars indicate the mean±SD. For each data set one representative of two similar experiments is shown.

In summary, serum antibody half-life was drastically shortened in chronically LCMV-infected mice, likely contributing to inefficient mAb-based lymphocyte subset depletion. This limitation was overcome by depletion-AAV technology. GC B cells, however, which are notoriously difficult to deplete irrespective of the infection context^74,75^, were partially resistant to AAV-αCD20-mediated depletion.

### Co-administered depletion-AAVs allow for the simultaneous depletion of two lymphocyte subsets

Next we set out to test whether depletion-AAVs might allow for the simultaneous depletion of two different lymphocyte subsets. For this we treated mice with AAV-ctrl, AAV-αCD20, AAV-αCD8 or with a combination of the latter two (combo), followed by acute LCMV infection (Fig. 5A). An assessment of spleens by flow cytometry (Fig. 5B-D) and immunohistochemistry (Fig. 5E) demonstrated that combo treatment depleted B cells and CD8 T cells equally efficiently as AAV-αCD20 and AAV-αCD8α single treatment, respectively.

**Figure 5:**
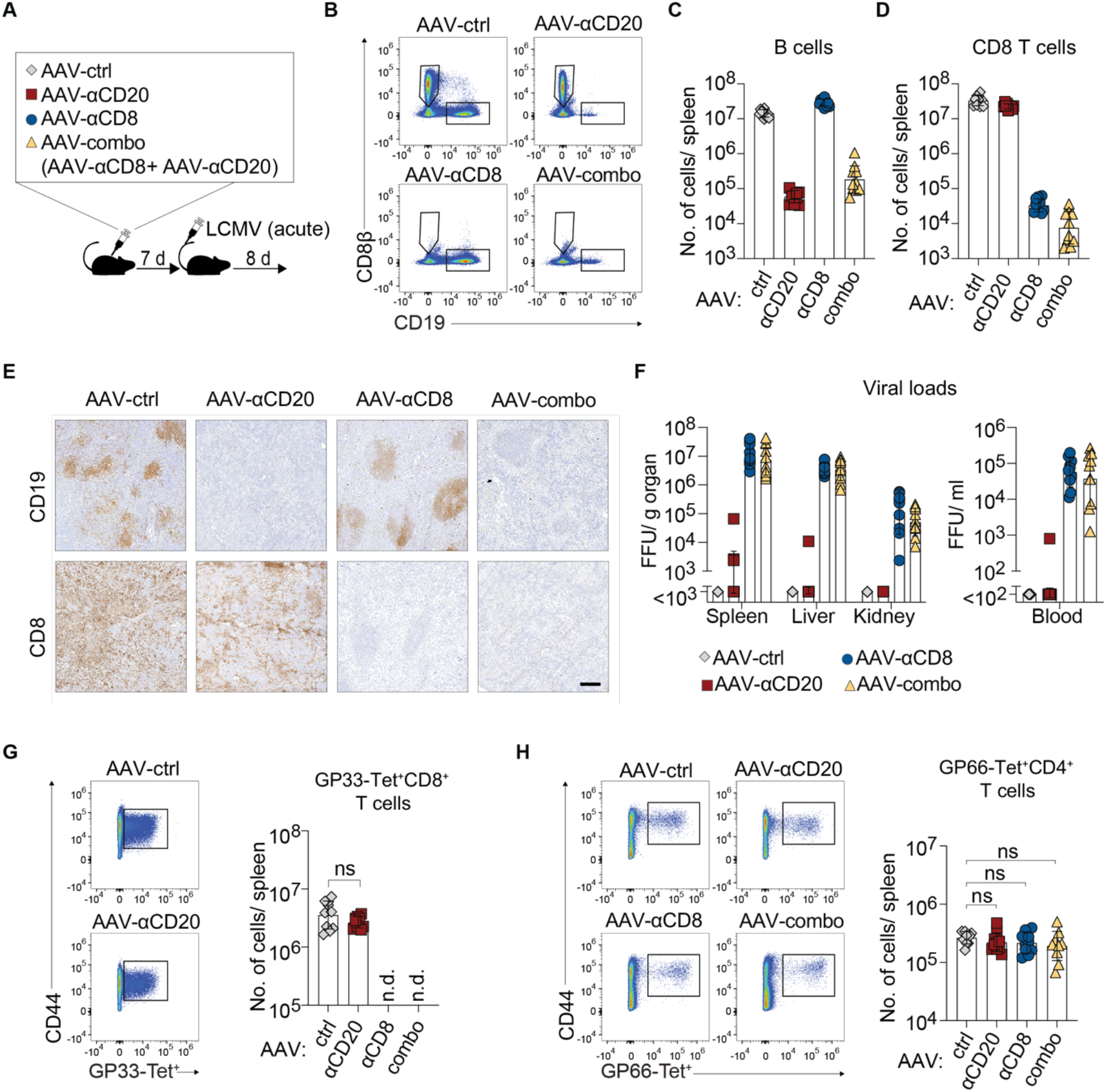
Co-administered depletion-AAVs allow for the simultaneous depletion of two lymphocyte subsets. We administered AAV-ctrl, AAV-αCD20, AAV-αCD8 or AAV-αCD20 in combination with AAV-αCD8 (combo) one week prior to LCMV infection (acute setting; A). Representative FACS plots (B; pre-gated on live lymphocytes) and absolute numbers of B cells (C) and CD8 T cells (D) in spleen on d8 after infection. Representative histological spleen sections from day 8 p.i. (E) stained for B cells (top row) or CD8 T cells (bottom row; magnification bar: 200 µm). Viral loads in spleen, liver, kidney and blood on d8 p.i. (F). Representative FACS plots of GP33-Tet^+^ CD8 T cells (pre-gated on CD4^−^CD8β^+^ live lymphocytes; left; G) and their total counts (right). Representative FACS plots of GP66-Tet^+^ CD4 T cells (pre-gated on CD4^+^CD8β^−^ live lymphocytes, left; H) and their total counts (right). Symbols in (C,D,F,G,H) represent individual mice and bars indicate the mean ± SD. (C,D,F,G,H) show combined data from two independent experiments, analyzed by unpaired t test (G) and one-way ANOVA followed by Dunnett’s post-test (H). ns: not statistically significant; n.d.: not determined.

In keeping with a central role for CD8 T cells in the clearance of acute LCMV infection^39–43^, mice treated with AAV-αCD8 or AAV-combo failed to clear LCMV from spleen, liver, kidney or blood (Fig. 5F), whereas AAV-ctrl-or AAV-αCD20-treated mice suppressed or controlled LCMV replication by day 9. Of note, AAV-αCD20-treated animals harbored normal numbers of GP33-specific CD8 T cells (Fig. 5G), confirming that B cells are dispensable for an efficient CD8 T cell response and control of acute LCMV infection^26,44,76–78^. Depletion of B cells or CD8 T cells or a combination thereof did not significantly affect the number of antiviral CD4 T cells reacting to the immunodominant GP-derived epitope GP66 either (Fig. 5H). In summary, the option to simultaneous deplete two or possibly even multiple lymphocyte subsets attests to the versatility of depletion-AAVs. Moreover, the present data corroborate a key role for CD8 T cells but not B cells in controlling acute LCMV infection and document that in this infection context neither one of the two is required for a substantial antiviral CD4 T cell response.

### Antiviral antibodies suppress chronic viremia to enable CD8 T cell-dependent clearance but B cells of irrelevant specificity promote virus-specific CD4 and CD8 T cell responses

The relative importance and interdependence of antibody-producing B cells and CD8 T cells in the control of chronic viral infection remains a subject of active investigation and controversial debate^25,41,44,46,47,49–51,79–84^. To address these questions, we treated mice with AAV-αCD20, AAV-αCD8 or AAV-ctrl or with a combination of the first two vectors (combo), followed by chronic LCMV infection one week later (Fig. 6A). A group of B cell-deficient *J_H_T* mice was included for comparison. AAV-ctrl-treated mice cleared viremia by day 82 whereas B cell-deficient *J_H_T* mice as well as B cell-depleted (AAV-αCD20) animals remained highly viremic for >100 days (Fig. 6B), corroborating a key role for B cells in the clearance of chronic viral infection. In contrast to B cell-depleted mice, CD8 T cell-depleted mice exhibited partial viral load control from around day 50 to day 80 after infection but subsequently experienced viral rebound^80^. Intriguingly, the depletion of CD8 T cells in addition to B cells did not result in higher viremia than B cell depletion alone. Taken together these findings indicated that B cell-dependent mechanisms afforded an initial suppression of viral loads. CD8 T cells were required to completely resolve residual viremia but failed to exert detectable antiviral effects when B cells were lacking. Animals treated with AAV-αCD20 and/or AAV-αCD8 were generally devoid of circulating B cells and/or CD8 T cells, respectively, throughout the experiment and the same was seen in spleen on day 112 (Fig. S5A-E). Of note, however, 1 out of 10 AAV-αCD8 single-treated mice, studied in two independent experiments, exhibited a restoration of the CD8 T compartment on day 112, despite efficient depletion at the beginning of the experiment. This was apparently due to an anti-idiotypic antibody response to the AAV-encoded CD8 antibody (Fig. S5F)^85–87^.

**Figure 6:**
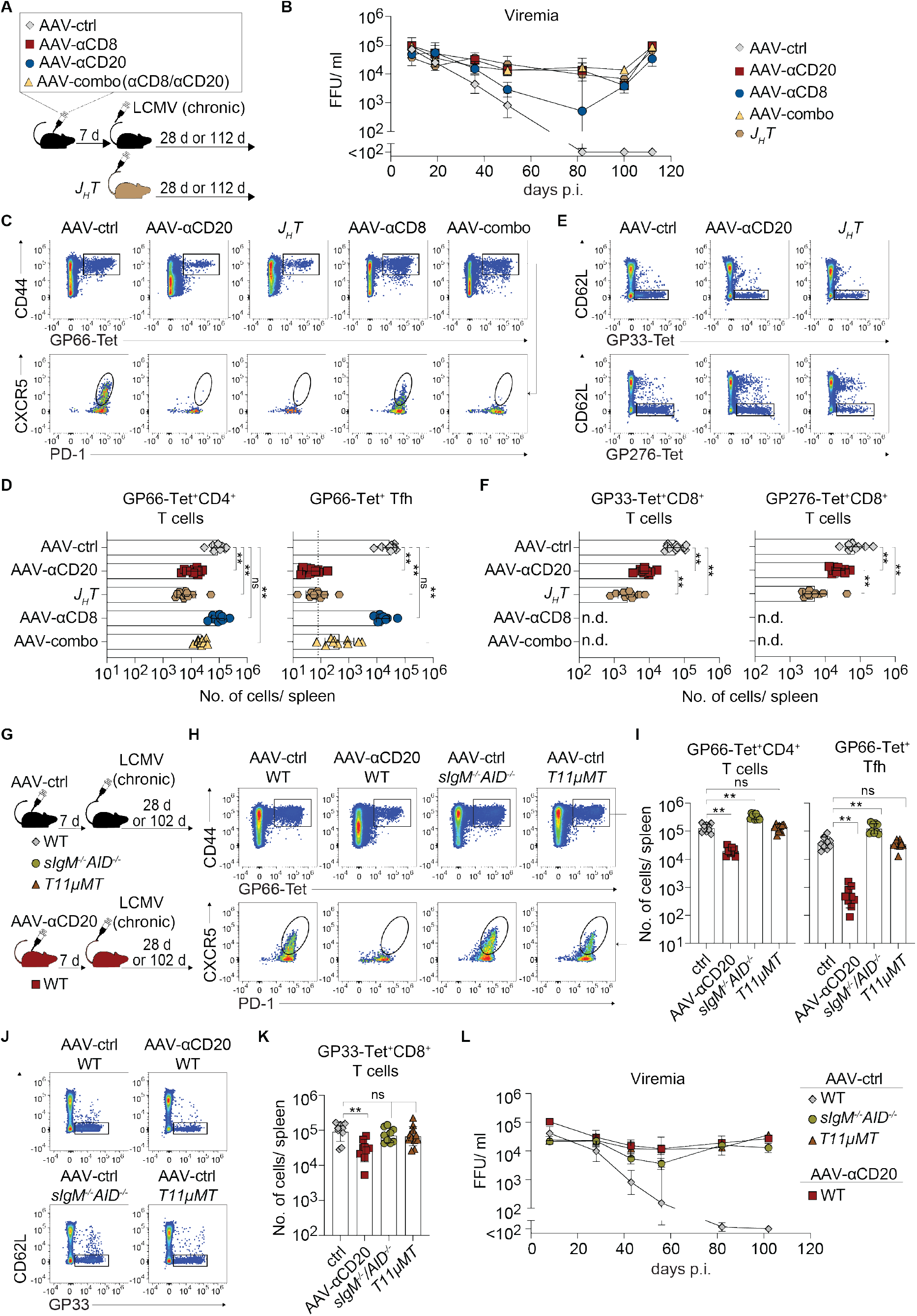
Antiviral antibodies suppress chronic viremia to enable CD8 T cell-dependent clearance but B cells of irrelevant specificity promote virus-specific CD4 and CD8 T cell responses. A-F: We administered AAV-ctrl, AAV-αCD20, AAV-αCD8 or AAV-αCD20 in combination with AAV-αCD8 (combo) to WT mice (A). One week later the treated WT mice and a group of untreated *J_H_T* mice were infected with LCMV (chronic). Viremia was monitored over time (B). Representative FACS plots (C; pre-gated on Ter119^−^CD4^+^ live lymphocytes) and absolute numbers of GP66-Tet^+^ CD4 T cells (D; left panel) and Tfh (D; right panel) in spleen on d28 after infection. Representative FACS plots (E; pre-gated on B220^−^CD4^−^Ter119^−^CD8β^+^ live lymphocytes) and absolute numbers of GP33-Tet^+^ CD8 T cells (F; left) and GP276-Tet^+^ CD8 T cells (F; right) on d28 p.i. in spleen. G-L: We administered AAV-ctrl to WT, *T11µMT* or *sIgM^−/−^AID^−/−^*mice and AAV-αCD20 to WT mice one week prior to chronic LCMV infection (G). Representative FACS plots (H; pre-gated on B220^−^CD8^−^CD4^+^ live lymphocytes) and absolute numbers of GP66-Tet^+^ CD4 T cells (I; left panel) and Tfh (I; right panel) on d28 p.i. in spleen. Representative FACS plots (J; pre-gated B220^−^CD4^−^Ter119^−^CD8β^+^ live lymphocytes) and absolute numbers of GP33-Tet^+^ CD8 T cells (K) on d28 p.i. in spleen. Viremia was monitored over time (L). Symbols in (B) represent the mean±SD of four (AAV-αCD8-treated group; M9 was excluded due to anti-drug antibodies, see Fig. S5D,F) to five mice per group (all other groups). One of two independent experiments is shown. Symbols in (D,F,I,K) represent individual mice and bars indicate the mean ± SD. Symbols in (L) represent the mean±SD of 4-5 mice. (D,F,I,K) show combined data from two independent experiments, analyzed by an ordinary one-way ANOVA followed by Dunnett’s post-test in (D,I,K) and Tukey’s post-test in (F). *: p<0.05; **: p <0.01; ns: not statistically significant; n.d.: not determined.

Marginal zone defects as observed in B cell-deficient GEMMs but not in AAV-αCD20-depleted mice (compare Fig. 3F) can negatively impact antigen presentation, thus potentially confounding earlier GEMM-based studies on the dependence of antiviral T cell responses on B cells^44–46,88^. To study the importance of B cells for T cell responses to chronic viral infection, an experiment was conducted analogously to the one above but analysis was performed on day 28 i.e. at a time point when viral loads in the different groups were still in similar ranges. (Fig. S5G). B cell-deficiency in *J_H_T* mice as well as in AAV-αCD20-depleted animals resulted in approximately 10- and 7-fold lower GP66-specific CD4 T cell responses, respectively, and caused a virtually complete absence of GP66-specific follicular T helper cells (Tfh; Fig. 6C-D)^74^. In contrast, the depletion of CD8 T cells affected neither total nor Tfh-differentiated GP66-specific CD4 T cells. B cell depletion resulted also in an 8-fold and 3-fold reduction in the number of GP33- and GP276-tetramer-binding CD8 T cells, respectively (Fig. 6E-F), which may have been a consequence of impaired CD4 T cell responses^89,90^. CD8 T cell responses were even further reduced in *J_H_T* mice, which was likely attributable to these animals’ defective splenic microarchitecture, attesting to the advantages of AAV-αCD20-mediated B cell depletion.

Impaired CD4 and CD8 T cell responses in AAV-αCD20-depleted mice raised the possibility that defective viral load control in such animals was due, at least in part, to indirect effects of B cell deficiency on T cell responses rather than to the lack of B cells and/or antiviral antibodies. To address this possibility, we studied T cell responses and viral load control in *sIgM^−/−^AID^−/−^* mice, harboring B cells with a diverse receptor repertoire but unable to secrete antibodies. In addition we studied *T11µMT* mice, which have a virtually monoclonal LCMV-unrelated B cell repertoire^91^ and thus are largely devoid of both LCMV-specific B cells and LCMV-specific antibodies. We compared T cell responses of chronically LCMV-infected *sIgM^−/−^AID^−/−^* and *T11µMT* mice (AAV-ctrl-treated) to those of either B cell-depleted (AAV-αCD20) or control-treated (AAV-ctrl) WT mice (Fig. 6G). When assessed on day 28 after infection, total and Tfh-differentiated GP66-specific CD4 T cells of *T11µMT* and *sIgM^−/−^AID^−/−^* mice were comparable or higher than in WT controls, whereas AAV-αCD20-mediated depletion suppressed these responses (Fig. 6H-I). Analogously, *sIgM^−/−^AID^−/−^* as well as *T11µMT* mice mounted GP33-specific CD8 T cell responses of normal magnitude (Fig. 6J-K). The number of CX3CR1^−^Ly108^+^ stem-like and CX3CR1^−^Ly108^−^ exhausted GP33-specific CD8 T cell subsets^90^ were unaffected in these animals whereas CX3CR1^+^Ly108^−^ effector cells were somewhat reduced, possibly as a consequence of incipient viral load differences (Fig. S6A,B). Importantly, however, and irrespective of their largely unimpaired CD4 and CD8 T cells responses, *sIgM^−/−^AID^−/−^* as well as *T11µMT* mice remained highly viremic and failed to suppress chronic LCMV viremia over an observation period of >100 days (Fig. 6L). Taken together these findings indicated that the mere presence of B cells, irrespective of their antigen-specificity, was essential for antiviral CD4 and CD8 T cells of normal magnitude. Clearance of chronic viremia, however, depended strictly on a concomitant antiviral antibody response.

## Discussion

This study establishes depletion-AAVs as an innovative approach to interrogate the role of individual lymphocyte subsets in various aspects of immunobiology. Moreover, AAV-mediated delivery of a decoy receptor for IL-33 allowed for the efficient blockade of IL-33 signaling to antiviral CD8 T cells in mice. As a somatic gene therapy approach depletion-AAVs combine key advantages of GEMMs with the versatility of the classical mAb-based method. Accordingly, key advantages of depletion-AAV technology over GEMMs comprise i) its functioning independently of mouse background, ii) the option to deplete more than one lymphocyte subset without the need for cross-breeding and, equally importantly, iii) the avoidance of compensatory mechanisms and accessory deficiencies that can result from the absence of lymphocyte subsets during ontogeny. Main advantages of depletion-AAVs over passively administered mAb depletion consists in the former’s permanent effect without the need for re-administration, avoiding also potentially confounding effects due to the re-emergence of only transiently mAb-depleted cells. Additionally, depletion-AAVs readily establish very high antibody concentrations (>500 µg/ml of serum), which is difficult to reach by passive mAb administration yet can be required for robust depletion under certain experimental conditions such as in persistent viral infection^65,66,68^.

However, depletion-AAVs have limitations, too, which notably comprise any potential off-target effects inherent to a vectorized mAb^92^. Furthermore, and as exemplified by the poor depletion of GC B cells in this study, the efficiency of antibody-mediated depletion can vary between tissue compartments. Unlike half-life-related limitations, this shortcoming of antibody-mediated depletion cannot be overcome by depletion-AAVs. A potential refinement of depletion-AAV technology allowing for the termination of depletion-mAb expression when needed, would further expand its utility and might facilitate the technology’s clinical exploitation.

Using depletion-AAVs as a methodological approach, the present findings corroborate and extend our understanding of the central role B cells play in chronic viral infection. More specifically our study shows that the importance of B cells extends well beyond their role as antibody producers^25^. We find that B cells are key to enable and/or sustain efficient antiviral CD4 and CD8 T cell responses to chronic viral intruders. B cells may affect CD8 T cell responses directly and/or indirectly. Given that CD8 T cell responses in chronic LCMV infection are known to depend on CD4 T cells and on Tfh cells in particular^33,41,89,90,93–96^, indirect B cell effects based on the promotion of CD4 T cell responses seem likely. At the same time, the positioning of stem-like CD8 T cells in the B zone^97–99^ is suggestive for additional, direct B cell effects on CD8 T cell responses. The experiments in *sIgM^−/−^AID^−/−^* and *T11µMT* mice indicate that B cell support for antiviral CD4 and CD8 T cell responses is independent of Fc receptor-dependent or BCR-mediated antigen uptake and presentation, respectively^100,101^. Moreover, AAV-αCD20 depletion shows that the aforementioned CD8 T cell-promoting effects of B cells extend beyond their essential role in the formation of the splenic microarchitecture.

Last but not least we observed that CD8 T cell depletion failed to result in a detectable increase in viral loads when B cells were depleted, too, indicating that the overall contribution of CD8 T cells in curbing high-level LCMV viremia is limited at best. In conjunction with transient viral load suppression and subsequent rebound viremia in CD8 T cell single-depleted animals these findings suggest antibody responses are the main initial player, suppressing viral loads to a certain threshold at which CD8 T cell responses take over to resolve viremia. Taken together the present study introduces depletion-AAVs as a versatile method to interrogate the role of lymphocyte subsets in immunobiology, offering key advantages over existing GEMM and mAb approaches. Exploitation of this novel tool offered new insights into the sophisticated interplay of B cells and T cells in the control of chronic viral infection, which should help to refine vaccination strategies for human diseases of global importance.

## Acknowledgments

We wish to thank Jean-Charles Paterna and the entire team of the University of Zurich Viral Vector Facility for vector batch production and technical advice, Lukas Jeker, Ondrej Stepanek, Camille Locht and Rolf Zinkernagel for providing hybridomas, Hartmut Hengel and Zsolt Ruzsics for MCMV, Cynthia Saadi and Min Lu for excellent technical assistance.

## Author contributions

A.L.K., A.-F.M., M.D., T.A.-M., Y.I.E., W.V.B., D.M. and D.D.P. designed experiments. A.L.K., A.-F.M., M.D., T.A.-M., Y.I.E., W.V.B., K.S., and I.W. performed experiments. A.L.K., M.K. and D.D.P. analyzed data. A.L.K. and D.D.P. wrote the manuscript.

## Material and Methods

### Animals and ethics statement

C57BL/6J WT mice were initially purchased from Charles River and were further bred at the Laboratory Animal Science Center (LASC) of the University of Zurich and at the ETH Phenomics Center (EPIC). BALB/c mice were purchased from Janvier Labs. *J_H_T* mice^102^, *T1-Fc^tg^*^103^, *IL-33^−/−^*^104^, *T11µMT*^105^ have been described and were bred at LASC and EPIC. *IL-33^−/−^* mice were generously provided by RIKEN under MTA, *T1-Fc^tg^* mice were obtained from the European Mouse Mutant Archive (EMMA). *sIgM^−/−^AID^−/−^* mice were obtained by cross-breeding *AID^−/−^*^106^ mice with *sIgM^−/−^* mice^107^ (Jackson Laboratories strain #003751). All mice were bred under specific pathogen-free (SPF) conditions. Within experiments mice were age- and sex-matched. However, both genders were used to reduce animal numbers bred for research purposes. All experiments were performed at the University of Basel in accordance with the Swiss law for animal protection and with authorization from the Cantonal veterinary office. Experimental groups were not randomized and the experiments were not conducted in a blinded fashion.

### Cell lines

We obtained NIH 3T3 cells from ATCC (CRL-1658) and cultured them in DMEM (Sigma, D0819; supplemented with 10 % fetal calf serum (FCS)). BHK21 cells were purchased from ECACC (85011433) and cultured in DMEM (supplemented with 10 % FCS, 10 mM HEPES (Gibco, 15630056), 1 mM Na-Pyruvate (Gibco, 11360070), 0.3 g/L tryptose phosphate broth (Sigma, T8782). FreeStyle 293-F suspension cells were obtained from Invitrogen/ ThermoFisher (R790-07) and cultured in HyClone CDM4HEK293 media (cytiva, SH30858.02; supplemented with 4 mM GlutaMax (Gibco, 35050038)). VeroE6 were purchased from ECACC and were cultured in in DMEM (supplemented with 10 % FCS). All cells were cultured at 37 °C in an atmosphere of 5 % CO_2_. All cell lines were tested for mycoplasma at regular intervals and were confirmed negative.

### Viruses, virus titrations and infections

We propagated lymphocytic choriomeningitis virus (LCMV) strain Armstrong Clone 13 (LCMV Cl13)^108^ on BHK-21 cells at a multiplicity of infection (MOI) of 0.01 and harvested virus-containing cell supernatant 48 h later. LCMV strain WE was originally obtained from Rolf Zinkernagel, University of Zurich, Switzerland. Recombinant LCMV strain Armstrong expressing the glycoprotein (GP) of strain WE (rARM/WE-GP)^109^ was grown on BHK21 cells at a MOI of 0.01 and harvested 68 h later. Recombinant LCMV expressing the GP of Vesicular Stomatitis Virus (VSVG; rLCMV/VSVG^61^) was propagated in FreeStyle 293-F suspension cells and virus containing supernatant was harvested 72 h later. Mouse cytomegalovirus (MCMV; MCMV BAC pSM3fr-MCK-2fl-M36GF) was generously provided by Hartmut Hengel and Zsolt Ruzsics, University of Freiburg, Germany. It was generated, propagated and titrated as described previously^110^. VSV Indiana was grown on BHK-21 cells at MOI of 0.01 and harvested 24 h later^111^.

We administered LCMV Cl13 intravenously (i.v.) into the tail vein at a dose of ≥ 2×10^6^ FFU resulting in a chronic infection. To induce an acute infection with LCMV, we infected mice with rARM/WE-GP at a dose of 2×10e^5^ FFU intraperitoneally (i.p.) in Fig. 1B-F, with rARM/WE at a dose of 200 FFU i.v. in Fig. 1G-I and with LCMV WE at a dose of 500 FFU i.v in Fig. 1J-L, Fig. 2 and Fig. 4. MCMV was administered i.p. at a dose of 1×10e^6^ PFU. rLCMV/VSVG was given i.v. at a dose of 6×10e^4^ PFU.

LCMV titers were determined by immunofocus assay following a protocol adapted from Battegay et al.^112^. In short, we prepared serial dilutions of virus stocks, viremia samples or organ samples in 96-well plates in MEM (Sigma, M0446; supplemented with 2 % FCS and 1 % Penicilline/ Streptomycin (P/S; ThermoFisher, 15140-122)). We added 3 x 10^4^ 3T3 cells to each well and incubated cells with virus for 2-3 h at 37 °C and 5 % CO_2_. After adding overlay (1 % methylcellulose (Sigma, M0430), 10 % FCS, 1 % P/S in DMEM), we incubated the cells for 48 h at 37 °C and 5 % CO_2_. Cells were fixed with 4 % paraformaldehyde (PFA; Sigma, 8187151000) in phosphate-buffered saline (PBS) for 10 min at room temperature (RT), followed by a permeabilization step with 1 % Triton-X (Sigma, T8787) in buffered saline solution (BSS) for 20 min at RT. Next, samples were blocked with 5 % FCS in PBS for 30 min, followed by incubation with the rat anti-LCMV-NP antibody VL4^112^ in PBS containing 2.5 % FCS for 1 h at RT. Plates were washed with tap water and then incubated with secondary horseradish peroxidase-(HRP-) conjugated goat anti-rat IgG antibody (Jackson, 112-035-003) diluted 1:500 in PBS containing 2.5 % FCS for 1 h at RT. Plates were washed and infectious foci were visualized by a colour reaction using 1.4 mM 3,3’-Diaminobenzidine tetrahydrochloride hydrate (Sigma, D-5637), 0.5 g/L Ammonium nickel(II)sulfate hexahydrate (Sigma, 09885) and 0.015 % H_2_O_2_ (Sigma, 216763) in PBS.

To determine LCMV viremia, we collected one drop of blood into 950 µl of BSS supplemented with 1 IE/ ml Heparin-Na (B. Braun, B01AB01) and analysed samples with the focus forming assay described above. To determine LCMV loads in organs, organ samples were harvested into MEM (supplemented with 2 % FCS and 1 % P/S) and weight was determined. Next, we homogenized organs with a stainless steel bead (Qiagen, 69989) using a tissue lyser (Qiagen). Samples were treated twice for 1 min at 30 Hz, centrifuged at >300 g for 5 min and analysed by focus forming assay.

To determine MCMV titers, we performed serial dilutions of organ samples, prepared analogously to LCMV organ samples as described above, in 96-well plates in MEM (supplemented with 2 % FCS and 1 % P/S). We added 3 x 10^4^ 3T3 cells to each well and incubated the cells for 2-3 h at 37 °C and 5 % CO_2_. After adding overlay (see above) we incubated the cells for 72 h at 37 °C in a 5 % CO_2_ atmosphere. Next, we fixed and permeabilized cells with ice-cold methanol (Sigma, 1060074000) for 20 min. After blocking with PBS supplemented with 5 % FCS for 10 min at RT, we incubated plates with anti-m123/IE1 antibody (clone IE1.01, generously provided by Stipan Jonjic, University of Rijeka, Croatia) for 1 h at RT. We washed plates and incubated them with a goat-anti-mouse-IgG secondary antibody coupled to HRP (Jackson, 115-035-062; 1:100) for 1 h at RT. Foci were visualized by means of a colour reaction performed as described above.

### Adeno-associated virus (AAV) vectors and administration

Plasmids for AAV vector production were generated by inserting the sequence of interest into the pENN.AAV.CB7.CI plasmid, which contains a CAG-promoter-driven expression cassette flanked by AAV2 inverted terminal repeats (ITRs), which was obtained from the PENN Vector Core (Perelman School of Medicine, University of Pennsylvania, PA, USA) under MTA. To allow antibody expression in a monocistronic manner from the AAV genome, we inserted an expression cassette consisting of building blocks assembled in the following order: a signal peptide for the heavy chain (HC), the HC variable domain (V_H_) of interest fused to the mouse IgG2a constant domain, a furin recognition site (R-K-R-R), a spacer (S-G-S-G), a F2A site for autocatalytic cleavage, a signal peptide for the light chain (LC), the LC variable domain (V_L_) of interest fused to the mouse kappa constant domain. The AAV-ST2-Fc construct was designed as follows: a signal peptide preceded the extracellular domain of ST2 (NCBI, Genbank: M24843.1), and the latter was coupled C-terminally to the mouse IgG1a constant domain. We used either one of two AAV vectors expressing antibodies of LCMV-unrelated, irrelevant specificity as controls, one encoding the VSVG-specific antibody VI10^54,91^ and the other encoding the mycobacterium-specific antibody E4H3^113^.The nucleotide sequences of the antibody variable regions of the E4H3 hybridoma (generously provided by Camille Locht, Institut Pasteur de Lille, France) as well as of the NK1.1-specific NK cell depletion antibody-producing hybridoma PK136 (generously provided by Ondrej Stepanek, Institute of Molecular Genetics of the Czech Academy of Sciences, Czech Republic) were determined by whole transcriptome shotgun sequencing (Absolute Antibody, UK). V_H_ and V_L_ of the αCD20-targeting antibody 18B12 and of the αCD8α-targeting antibody YTS169.4 were extracted from patents US 2007/0136826 A1 and WO2014025828 A1, respectively^114,115^. V_H_ and V_L_ sequences of the αCD4-targeting antibody hybridoma YTS191 and of the αGr1-antibody-producing hybridoma RB6.8C5 were sequenced as described below. Antibody expression cassettes were synthesized by Genscript and were subcloned into pENN.AAV.CB7.CI for vectorization. AAV vector production (in an AAV8 capsid format) and titration was performed by the Viral Vector Facility (VVF) of the Neuroscience Center Zurich (ZNZ), Switzerland. AAV vectors were administered to mice intramuscularly at doses of 2 x 10^10^ to 1 x 10^11^ viral genomes per mouse.

### Hybridoma sequencing, monoclonal antibody production and *in vivo* administration

To determine V_H_ and V_L_ sequences of the CD4-targeting antibody YTS191 and of the Gr1-targeting antibody RB6.8C5 (generously provided by Rolf Zinkernagel, University of Zurich, Switzerland, and by Lukas Jeker, University of Basel, Switzerland), we cultivated the corresponding rat hybridomas in IMDM (Gibco, 12440-053; supplemented with 10 % FCS, 1X MEM Non-essential amino acid solution (ThermoFisher, 11140050), 2 mM Glutamax (ThermoFisher, 35050061), 50 µM β-mercaptoethanol (Sigma, 8.05740) and 1 % P/S) and isolated cellular RNA using Direct-zol^TM^ RNA MicroPrep (ZymoResearch, R2062). To identify the V_H_, we designed reverse primers binding to both, rat-IgG2b (NCBI, GenBankM28671.1) and rat-IgG2a (GenBank M28669.1; 5’-TTGTCCACCTTGGTGCTGCT-3’; 5’-TCACTGAGCTGGTGAGAGTGT-3’; 5’-GGTGACTGGCTCAGGGAAA-3’). For sequencing V_L_, we designed reverse primers binding to the rat Ig kappa constant domain (GenBank: DQ402471.1; 5’-CACTCATTCCTGTTGAAGCT-3’; 5’-CAGGTATAGAGGTTATGCCTTTCA-3’; 5’-CTGATGTCTCTGGGATAGAAGT-3’). To circumvent challenges related to the unknown 5’ end of the respective mRNAs, precluding the design of a sequence-specific forward primer at the beginning or upstream of the V_H_ and V_L_, we adapted the strategy published by Meyer et al.^116^. We made use of the SMART (switching mechanism at 5’end of RNA transcript) technology. In short, cDNA libraries with the above-mentioned primers were generated using the Moloney murine leukemia virus (MMLV) reverse transcriptase, which adds several deoxycytosines to the 3’end of the cDNA transcript. Next, a primer (“template-switch oligo”; 5’-AAGCAGTGGTATCAACGCAGAGTACATrGrGrG-3’), which consists of a sequence of interest followed by three riboguanines, pairing with the cDNA deoxycytosine overhang of the cDNA and resulting in a template switch of the MMLV reverse transcriptase, extending the 3’ end of the cDNA by the sequence of interest. This creates a known, specific sequence at the cDNA 3’ end, which serves as template sequence for a forward primer in the PCR reaction (universal fwd primer). For the cDNA reaction, we mixed 1 µl of RNA (ca. 50 ng/µl) with 1 µl of 10 µM reverse primer and 1 µl of 10 mM dNTPs and incubated the mix for 3 min at 72 °C to denature any RNA secondary structures. We used two different heavy chain reverse primers and two different light chain primers in separate reactions for each hybridoma. Next, we mixed 2 µl of 5X SMART buffer, 1 µl 20 mM DTT, 2.7 µl ddH_2_O, 0.3 µl of 100 µM “template-switch oligo” primer, 0.5 µl of 40 U/ µl RNase Inhibitor and 0.5 µl of the SMARTScribe RT (Takara, 639537), added the mix to the RNA mix and incubated the reaction at 42 °C for 60 min, followed by 5 min at 70 °C to stop cDNA synthesis. Next, we amplified the cDNA via polymerase chain reaction (PCR). For this we mixed 3 µl of the obtained cDNA with 8 µl 5X Phusion buffer, 3.2 µl of 2.5 mM dNTPs, 0.4 µl of 2000 U/ml Phusion® High-Fidelity DNA Polymerase (NEB, #M0530S), 2 µl of the universal forward primer (5’-AAGCAGTGGTATCAACGCAGAG-3’) and 2 µl of a reverse primer located upstream of the reverse primer used for cDNA synthesis (5’-TCACTGAGCTGGTGAGAGTGT-3’ or 5’-GGTGACTGGCTCAGGGAAA-3’ for V_H_ and 5’-CAGGTATAGAGGTTATGCCTTTCA-3’ or 5’-CTGATGTCTCTGGGATAGAAGT-3’ for V_L_) to perform a nested PCR. Cycling conditions were: 98 °C 30 s, then 35 cycles of: 98 °C 15 s, primer annealing for 30 s (primer-specific temperatures) and amplification at 72 °C for 30 s; final extension at 72 °C for 7 min). PCR products were purified from 2 % agarose-gels using QIAquick Gel Extraction Kit (QIAGEN, 28706) for Sanger sequencing (Microsynth).

For the YTS191 antibody we identified mixed sequencing signals for the light chain, suggesting the existence of a second light chain. To identify both light chains we performed subcloning of the light chain PCR products into pGEM-T-Vector System (Promega, A3600). First, we performed an A-tailing of the PCR products of the YTS191-light chain cDNA by adding 5 µl of 10X ThermoPol Buffer (NEB, B9004), 10 µl of 10 mM ATP, 0.2 µl of *Taq* DNA Polymerase (NEB, M0267S) to 30 µl of PCR reaction, filled up to 50 µl with ddH_2_O and incubated at 72 °C for 20 min. Next, we purified the A-tailed PCR product with a QIAquick PCR Purification Kit (QIAGEN, 28104), followed by subcloning into the pGEM-T-Vector according to the manufactures protocol. V_L_ sequences were identified by Sanger Sequencing from the transformed bacterial colonies (Microsynth).

Identified antibody sequences were synthesized by Genscript and subcloned into mammalian expression vectors for recombinant expression in an IgG2a format as described^117^. In brief, monoclonal antibodies were obtained by co-transfection of Expi-CHO cells with the HC- and LC-encoding plasmids of the corresponding antibody (Protein Expression Core Facility, PECF, of the Swiss Federal Technical Highschool, EPFL, Lausanne, Switzerland). Next, we purified antibodies from cell supernatant on an Äkta Primeplus purification system using HiTrap Protein G columns (cytiva, 17040401). We dialysed purified antibodies (Sigma, D9277) in PBS for 24 h at 4 °C and 1 h at RT. Antibodies were sterile-filtered and quantified with Pierce™ BCA Protein Assay Kits (Thermo Scientific, 23227). For antibodies PK136, E4H3 and YTS191 we identified two different light chains. Therefore, we expressed both LCs separately in combination with the corresponding HC and determined the correct light chain by testing the biological activity of the resulting constructs. To deplete CD4 T cells by recombinant antibody, we injected the purified monoclonal antibody YTS191-IgG2a twice i.p. at a dose of 200 µg on day −3 and −1 of infection. For the determination of antibody half-life during chronic LCMV infection, we injected 2 mg of E4H3-IgG2a intravenously 19 days after LCMV infection.

### Flow cytometry

To obtain a single cell suspension of splenocytes we mechanically disrupted spleens using a metal grid and resuspended splenocytes in FACS media (RPMI (Sigma, R2405) supplemented with 20 mM HEPES, 5 mM MgCl_2_ (ThermoFisher, AM9530G), 1 mM Na-Pyruvate, 1X MEM non-essential amino acids (Sigma, M7145), 5 % FCS, 1 % P/S, 0.05 mM β-mercapto-ethanol (Sigma, 8.05740)). All media and buffers were adjusted to mouse osmolarity^118^. Lymph nodes were mechanically disrupted using a 70 µm cell strainer followed by resuspension in FACS media. To analyze bone marrow we harvested the tibia, flushed the bone with FACS media to isolate the marrow and obtained a single cell suspension by passing cells through a 70 µm cell strainer.

Prior to staining, samples were incubated for 10 min at RT with DNAse I (50 µg/ml; Roche, 10104159001) and Dasatinib (50 nM; Sigma, CDS023389) in FACS media. All stains were performed in staining buffer (PBS supplemented with 10 % Opti-MEM, 1 % FCS, 1 % P/S, 10 mM HEPES, 5 mM MgCl_2_, 1 mM Na-Pyruvate, 1x MEM non-essential amino acids, 0.05 mM β-mercaptoethanol, 1 g/ ml D-Glucose (Gibco, A2494001)) containing rat IgG (Sigma, I4131) and anti-mouse CD16/CD32 (2.4G2; BioXcell, BE0307) to block unspecific antibody binding. BD Horizon^TM^ brilliant stain buffer (BD, 566385) was used whenever more than one brilliant violet dye was in the staining mix. Antibodies against B220 (RA3-6B2), Bcl-6 (K112-19), CD101 (Moushi101), CD11b (M1/70), CD11c (N418), CD127 (A7R34), CD138 (281-2), CD162 (2PH1), CD19 (1D3, 6D5), CD21 (7G6), CD23 (B3B4), CD25 (PC61), CD3 (145-2C11), CD38 (90, REA616), CD4 (RM4-5), CD44 (IM7), CD45.2 (104), CD49b (HMα2), CD62L (MEL-14), CD69 (H1.2F3), CD8α (53-6.7), CD8β (53-5.8), CX3CR1 (SA011F11), CXCR5 (L138D7), CXCR6 (SA051D1), FoxP3 (MF-14), GL7 antigen (GL7), IgD (11-26c.2a), IgM (IL/41, REA979), KLRG1 (2F1/KLRG1), Lag3 (C9B7W), Ly-108 (13G3), Ly-6C (AL-21, HK1.4), Ly-6G (1A8), NK1.1 (PK136), NKp46 (29A1.4), PD-1 (29F.1A12), Siglec H (551), T-bet (o4-46), TACI (8F10), Tcf-1 (C63D9), Ter119 (TER-119), Tim-3 (5D12/TIM-3) were used for staining and purchased from BD, Milteny, Biolegend, Cell Signaling and ThermoFisher (ebiosciences, Invitrogen).

MHC class II tetramers (I-A^b^) loaded with the GP_66-77_ peptide (DIYKGVYQFKSV) were obtained from the NIH Tetramer core facility and were used to identify LCMV-GP specific CD4 T cell responses. Tetramer staining was performed in combination with staining for chemokine receptors CXCR5 and CXCR6 for 1 h at RT in staining buffer supplemented with Dasatinib (50 nM) and biotin (20 nM; Sigma, B4501). After washing of the samples with FACS buffer (PBS supplemented with 1 % FCS), surface staining was performed for 30 min at RT.

To detect LCMV specific CD8 T cell responses in C57BL/6 mice we used MHC class I tetramers (H-2D^b^) loaded with the GP_33-41_ peptide (KAVYNFATC), with the GP_276-286_ peptide (SGVENPGGYCL) or with the NP_396-404_ peptide (FQPQNGQFI). To detect LCMV-NP specific CD8^+^ T cell responses BALB/c mice, we used MHC class I tetramers (H-2L^d^) loaded with the NP_118-136_ peptide (RPQASGVYM). MHC class I tetramers were obtained from the NIH Tetramer core facility or from the Tetramer core facility of the University of Lausanne. MHC class I tetramers were stained in combination with surface staining for 30 min at RT. Subsequent to tetramer and surface antibody staining, cells were washed with PBS and stained for dead cells using Zombie UV Fixable Viability Kit (Biolegend, 423108) or Zombie NIR Fixable Viability Kit (Biolegend, 423105) at a dilution of 1:2000 or 1:3000, respectively, for 15 min at RT. Samples were washed with FACS buffer and fixed in 2 % PFA for 10 min at RT. Samples were washed once more prior to acquisition.

We used the eBioscience™ Foxp3 / Transcription Factor Staining Buffer Set (00-5523-00, Invitrogen) to stain for transcription factors. After tetramer, surface and live/dead staining we prepared the Foxp3 Fixation/Permeabilization working solution and fixed splenocytes for 1 h. After washing twice with the 1X working solution of Permeabilization Buffer, we stained for transcription factors at 4 °C for 12 – 16 h. Samples were washed twice with 1X Permeabilization Buffer before acquisition.

For staining of cells in peripheral blood, 2-3 droplets of blood were collected into blood FACS buffer (PBS supplemented with 2 % FCS, 5 mM EDTA and 0.05 % sodium azide). Cells were stained for 20 min at 4 °C. After washing the cells with blood FACS buffer we performed red blood cell lysis and fixation using BD FACS™ Lysing Solution 10X Concentrate (BD, 349202) diluted 1:10 in ddH_2_O, incubated for 5 min at RT and washed twice before acquisition.

Samples were acquired on a BD LSRFortessa flow cytometer (BectonDickinson) and on a 5-laser AURORA spectral flow cytometer (Cytek Biosciences), followed by analysis using FlowJo (BectonDickinson) and Spectroflo (Cytek Biosciences) software.

### Enzyme-linked immunosorbent assay and neutralization assay

To determine antibody responses in serum we conducted an enzyme-linked immunosorbent assay (ELISA). For this we coated 96-well high-affinity binding plates overnight at 4 °C with coating antibody diluted in coating buffer (15 mM Na_2_CO_3_ and 35 mM NaHCO_3_ in ddH_2_O, pH: 9.6). After flicking off the coating mix, we added 5 % milk in PBS supplemented with 0.05 % Tween-20 (PBST; Sigma, P9416) for 1 h at RT to block non-specific binding. In a separate plate we prepared 3-fold serial dilutions of serum samples in PBST supplemented with 1 % FCS, transferred the diluted samples into the coated and blocked ELISA plates and incubated the samples for 1 h at 37 °C shaking at 300 rpm. Next, plates were washed 3 times with PBST and incubated with a secondary HRP-coupled antibody in PBST supplemented with 1 % FCS for 1 h at RT at 300 rpm. Plates were washed 3 times with PBST and once with PBS prior to adding the colour reaction mix (0.5 mg/ ml ABTS (Thermo Scientific, 34026), 28 mM citric acid, 44 mM (Na_2_HPO_4_) and 0.1 % H_2_O_2_ in ddH_2_O), which was incubated for 15-30 min. We terminated the colour reaction by adding 1 % of SDS (Sigma, 71729) in ddH_2_O and measured optical density at a wavelength of 405 nm on an Infinite® M Plex device (TECAN).

To detect (exogenous) IgG2a in the serum of C57BL/6 WT mice we used 1 µg/ml of a goat anti-mouse IgG2a (Jackson, 115-005-206) for coating and a goat anti-mouse IgG-HRP as secondary antibody (Jackson, 115-035-062; diluted 1:5000 for use). To identify LCMV nucleoprotein (NP)-specific antibodies, we coated plates with 1 µg/ml of recombinant NP^119^ and used a goat anti-mouse IgG-HRP (γ-chain specific) as secondary antibody (Sigma, A3673-1ML; diluted 1:1000). To measure anti-drug antibody responses directed against the AAV-delivered YTS169-IgG2a we coated plates with 1 µg/ml of YTS169-IgG2a or YTS191-IgG2a and used a goat anti-mouse IgG1-HRP (Jackson, 115-035-205; diluted 1:5000) as secondary antibody.

To determine absolute antibody concentrations of passively transferred or AAV-vectored antibodies we generated standard curves using a monoclonal antibody of identical isotype and intrapolated the OD values obtained from experimental serum samples. To quantify host-endogenous antibody responses we calculated the serum dilution, which reached half-maxium OD (EC_50_). We used naïve serum to determine the background of the assay.

In order to calculate antibody half-life (*t*_1/2_) of passively administered mAb we used a one-phase decay model^73^. To distinguish between the initial rapid distribution phase and the subsequent elimination phase of the antibody we excluded values of the early time points (2 h, 12 h and 24 h) from the calculation of antibody half-life ^73,120^.

To determine VSVG-specific antibody titers in serum we performed virus neutralization assays as previously described ^111^. Pre-diluted samples were heat-inactivated for 30 min at 56 °C. Next, we performed serial dilutions of samples in 96-well U-bottom plates in MEM supplemented with 2 % FCS. Samples were incubated with approximately 50 PFU of VSV-IND^111^ for 90 min at 37 °C. Next, we transferred the serum – virus mixtures into a 96-well flat-bottom plate and added 3 x 10^4^ VeroE6 cells per well. After an incubation of 1 hour at 37 °C we added 80 µl of overlay (1 % methylcellulose, 10 % FCS, 1 % P/S in DMEM). We incubated the assay for 16-18 h at 37 °C before fixing and staining with crystal violet solution (0.5 % crystal violet, 48 % ethanol, 1.85 % formalin, 0.8 % NaCl in ddH_2_O). We counted PFU in each well and calculated the dilution resulting in 50 % neutralization (IC_50_).

### Immunohistochemistry

We performed histological analyses on snap-frozen tissue. For CD19 (6OMP31; ebioscience, 14-0194-80), F4/80 (BM8; Biolegend, 123108), CD8α (4SM16; Invitrogen, 14-0195-82) and CD169 (MOMA-1; Biorad, MCA947GA) immunohistochemistry, 5 µm cryosections were fixed in 2 % PFA for 15 min. To avoid unspecific binding, sections were incubated successively with a peroxidase-blocking solution (Dako, S2023), Fab fragments of anti-mouse IgG (Jackson ImmunoResearch, 115-007-003) and 2.5 % goat serum (Vector Laboratories, S-100). Slides were incubated with the primary antibody overnight and specific bonds were visualized using HRP conjugated secondary antibody against rat IgG and polymerized 3,3ʹ-diaminobenzidine (Dako, K3468). Nuclei were counterstained with Mayer’s Hemalum solution (Merck 1.09249.05).

For SIGN-R1 (ERTR9; BMA biomedical, T-2024) immunohistochemistry, 5 µm cryosections were fixed in 1% PFA for 5 min. To avoid unspecific binding, sections were incubated successively with a peroxidase-blocking solution (Dako, S2023), avidin/biotine block solution (Biolegend, 750001835/36) and 10 % mouse serum (Jackson ImmunoResearch, 015-000-120). Slides were incubated with the primary antibody overnight and specific bonds were visualized using an HRP conjugated streptavidin and polymerized 3,3ʹ-diaminobenzidine (Dako, K3468). Nuclei were counterstained with Mayer’s Hemalum solution (Merck, 1.09249.05)

### Quantification and statistical analysis

For statistical testing we used GraphPad Prism software (Version 10.2.3, Graph Pad Software). When comparing two groups, we used an unpaired two-tailed Student’s t test. When more than two groups were compared One-Way ANOVA was performed, Dunnett’s post-test served to compare all groups against one reference group whereas Tukey’s post-test was conducted to compare all groups against each other. P values p<0.05 were considered statistically significant (*), p<0.01 as highly significant (**) and p>0.05 as not statistically significant. Absolute numbers were log-converted to obtain a near-normal distribution for statistical analysis.

---

**Figure S1:**
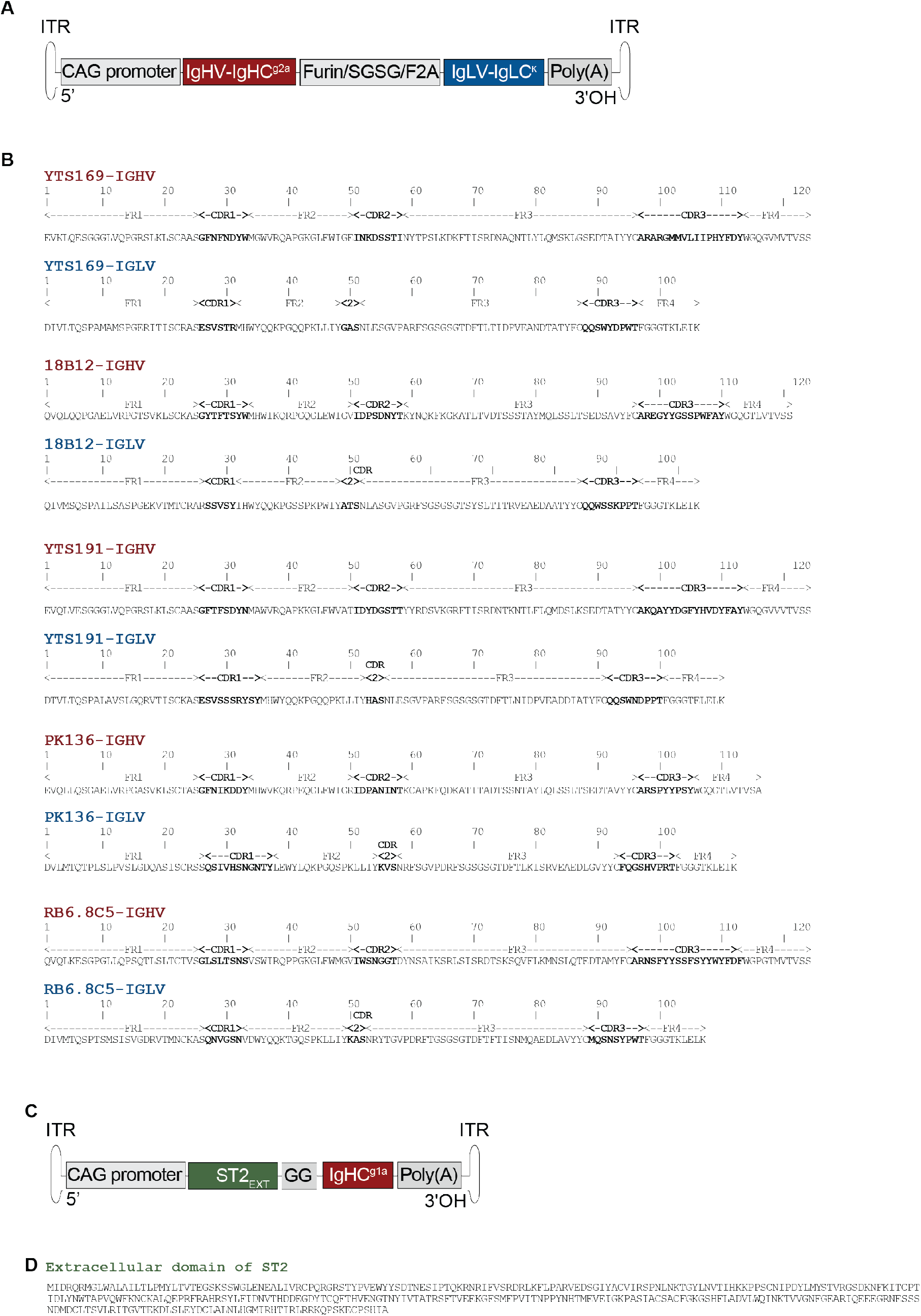
Genetic design and transgene sequences of various depletion-AAVs and of AAV-ST2-Fc. A: Schematic of the AAV-genome used to deliver an expression cassette for various depletion antibodies. ITR: inverted terminal repeats; CAG: chicken gamma actin; IgHV-IgHC^g2a^: variable domain of the heavy chain and constant mouse IgG2a domain; SGSG: serin-glycine-linker; Furin: furin cleavage site; F2A: Foot and mouth disease virus 2A peptide; IgLV-IgLC^ĸ^: variable domain of the light chain and kappa light chain constant domain; poly(A): polyadenylation signal. B: Amino acid sequences of variable regions of heavy (IGHV; red) and light chains (IGLV; blue) of depletion antibodies used in this manuscript. Framework regions (FR) and complementarity-determining regions (CDR; bold) were annotated according to IMGT (IMGT/DomainGapAlign). C: Schematic of the AAV-ST2-Fc construct. EXT: extracellular; GG: glycine-linker; IgHC^g1a^: mouse IgG1 constant domain. D: Amino acid sequence of the ST2 extracellular domain.

**Figure S2:**
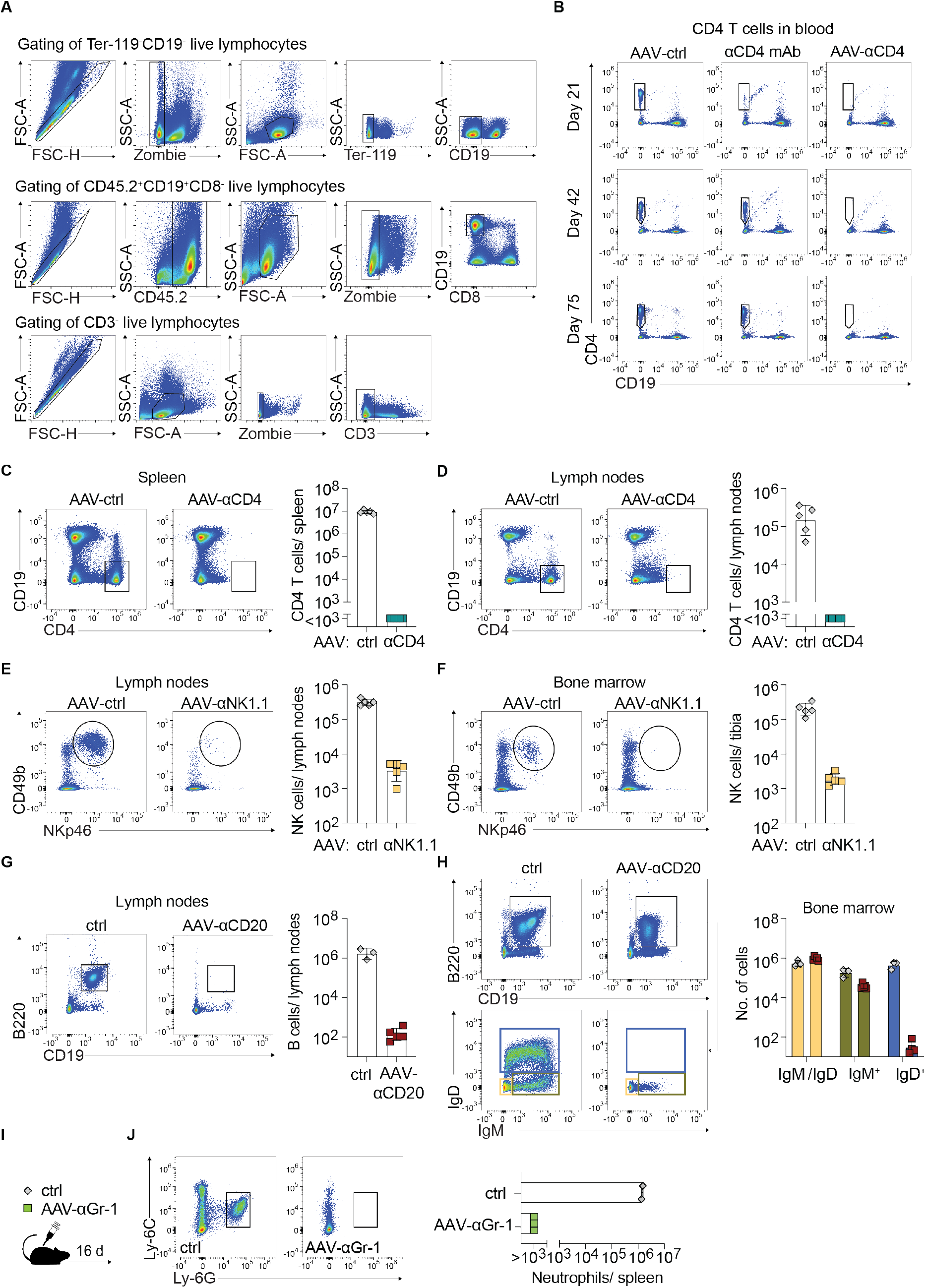
Depletion-AAVs in lymph nodes and bone marrow. A: Gating strategies of different lymphocyte populations. B: Mice were treated with AAV-αCD4 or AAV-ctrl one week prior to acute LCMV infection or were administered anti-CD4 depletion antibody (αCD4 mAb) as recombinant protein on d-3 and d-1 of infection. Representative FACS plots of CD4 T cells (pre-gated on lymphocytes) in blood on d21, d42 and d75 after LCMV infection as quantified in Fig.1C. C-D: Representative FACS plots and absolute numbers of CD4 T cells in spleen (C) and lymph nodes (D) 31 days after treatment with AAV-ctrl or AAV-αCD4 (same experiment as reported on in Fig. 1G; pre-gated on CD45.2^+^ live lymphocytes). E-F: Representative FACS plots and absolute numbers of NK cells in lymph nodes (E) and bone marrow (F) 31 days after treatment with AAV-ctrl or AAV-αNK1.1 (same experiment as reported on in Fig. 1M; pre-gated on CD3^−^ live lymphocytes). G-H: Representative FACS plots and absolute numbers of B cells in lymph nodes (G) 90 days after treatment with AAV-αCD20 (same experiment as reported on in Fig. 1P), side-by-side with untreated control mice (pre-gated on live lymphocytes). Representative FACS plots of B cells in bone marrow (H, top row; pre-gated on CD3^−^ live lymphocytes) and B cell subsets (G, bottom row), as well as their quantification (H, right). I-J: We treated mice with AAV-αGr-1 or left them untreated. Representative FACS plots and absolute numbers of neutrophils in spleen (J) on day 16 after treatment (pre-gated on CD45.2^+^CD3^−^CD19^−^ live lymphocytes). Symbols in (C-H) and (J) represent individual mice, bars indicate the mean±SD. Panels A-H show one representative of two independent experiments.

**Figure S3:**
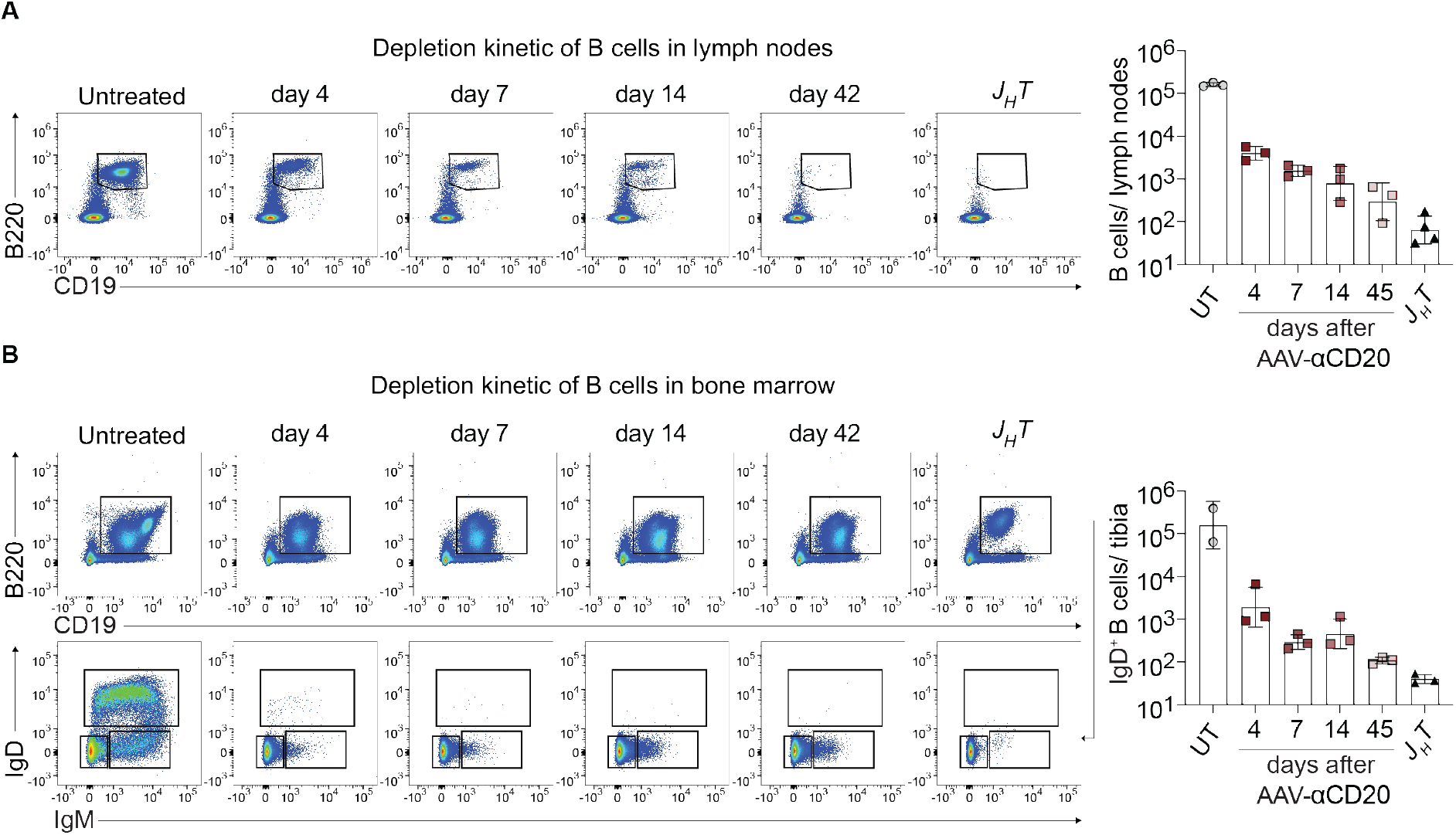
Kinetics of depletion-AAV-mediated B cell elimination in lymph nodes and bone marrow. We administered AAV-αCD20 to mice and monitored the depletion of B cells in lymph nodes (A) and bone marrow (B) over time (same experiment as reported on in Fig. 3A). Untreated (UT) WT and *J_H_T* mice are shown for comparison. Representative FACS plots and absolute numbers of B cells in lymph nodes at the indicated time points (A; pre-gated on live lymphocytes). Representative FACS plots of B cells (B, top row; pre-gated on CD8^−^ live lymphocytes) and B cell developmental stages (B; bottom row, identified by IgD and IgM expression) in bone marrow, as well as the quantification of mature B cells (IgD^+^). Symbols represent individual mice and bars indicate the mean±SD. For each data set one representative of two independent experiments is shown.

**Figure S4:**
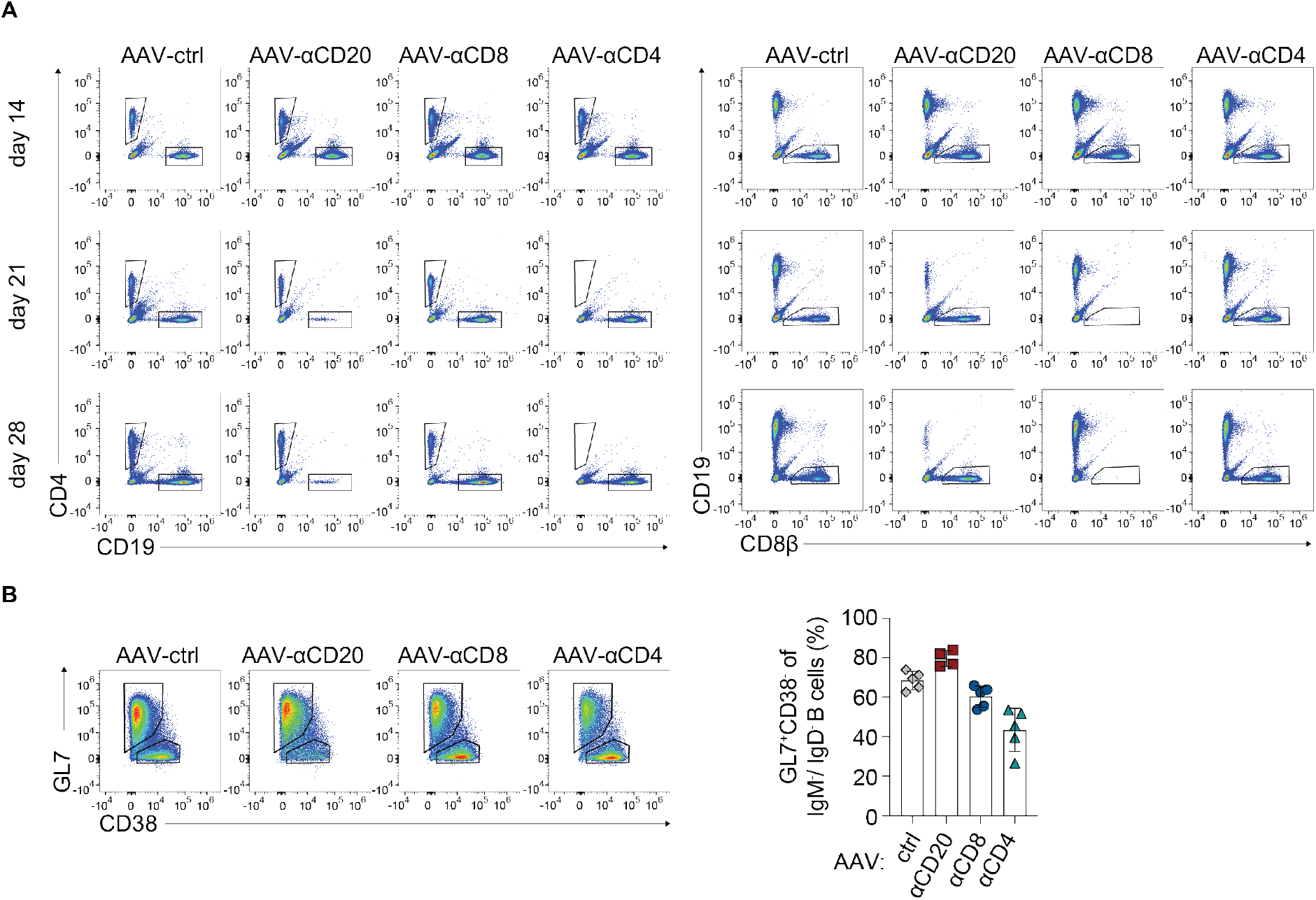
Depletion-AAVs are effective during chronic LCMV infection. Exemplary FACS plots of CD8 T cells, CD4 T cells and B cells (pre-gated on lymphocytes) in blood on day 14, 21 and 28 after LCMV infection (chronic) from mice treated with AAV-ctrl, AAV-αCD20, AAV-αCD8 or AAV-αCD4 14 days after infection as shown in Fig. 4C,D (A). Representative FACS plots of GC (GL7^+^CD38^−^) B cells pre-gated on class-switched B cells (IgM^−^ IgD^−^; Fig. 4G) and their frequencies in spleen on d28 p.i. (B).

**Figure S5:**
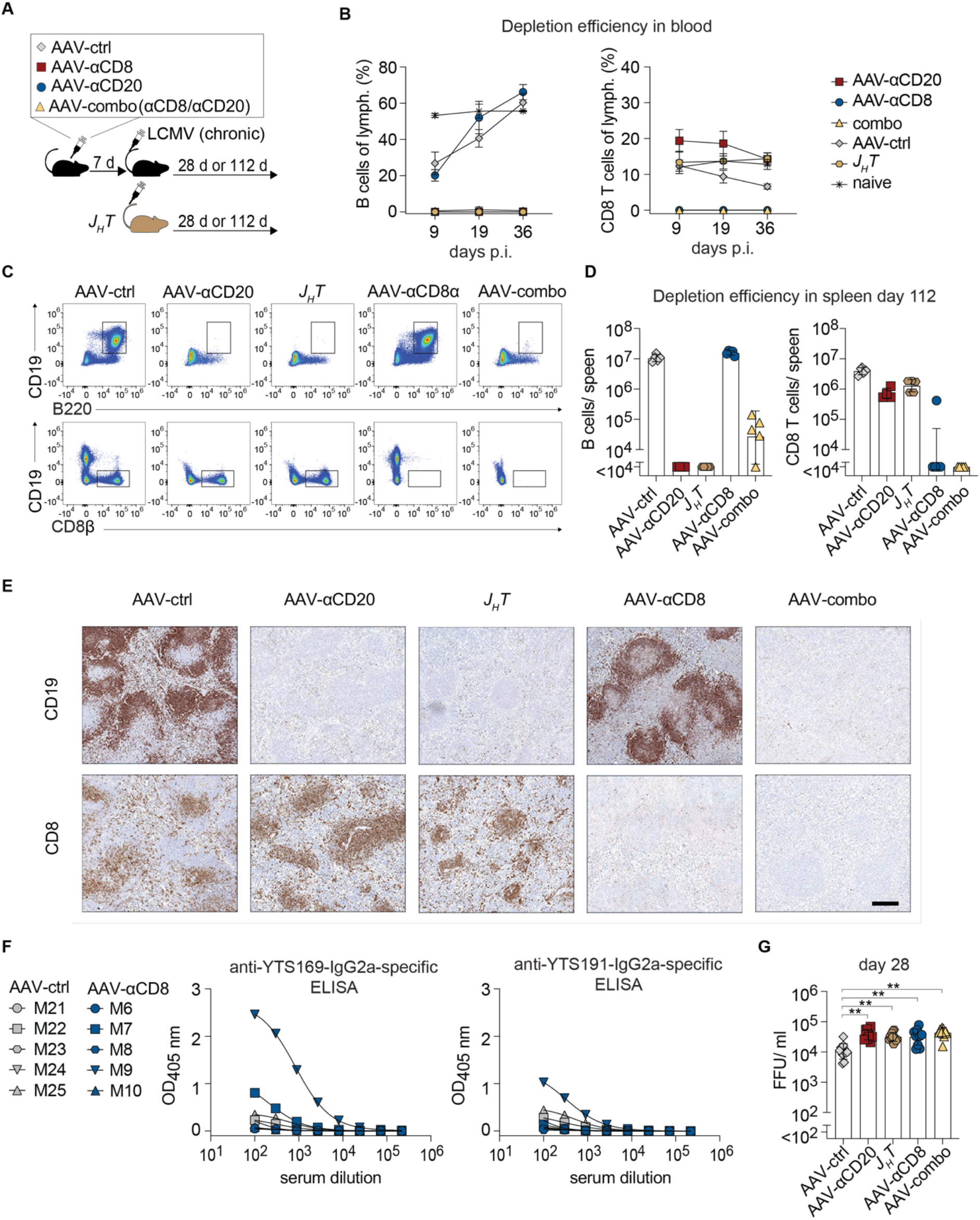
Efficiency of depletion-AAV-mediated B cell and CD8 T cell elimination in chronic LCMV infection. We administered AAV-ctrl, AAV-αCD20, AAV-αCD8 or AAV-αCD20 in combination with AAV-αCD8 (combo) to WT mice (A; same experiment as reported on in Fig. 6A,B). One week later the treated WT mice and an untreated control group of *J_H_T* mice were infected with LCMV (chronic). Percentages of B cells (B; left) and CD8 T cells (B; right) amongst peripheral blood lymphocytes. Representative FACS plots (C) of B cells (top row; pre-gated on CD4^−^CD8^−^Ter119^−^ CD45.2^+^ live lymphocytes) and CD8 T cells (bottom row; pre-gated on Ter119^−^ live lymphocytes) in spleen on day 112 after infection, as well as their quantification (D). Representative histological spleen sections from day 112 p.i. (E) were stained for B cells (top row) or CD8 T cells (bottom row; magnification bar: 200 µm). From AAV-ctrl- and AAV-αCD8-treated mice in the experiment to Fig. 6A,B we determined the binding of d112 serum antibodies to YTS169-IgG2a (F; left) and to YTS191-IgG2a (right). Viremia on day 28 p.i. in the experiment reported in Fig. 6C-F is shown in (G). Symbols in (B) represent the mean±SD of five mice per group. In panels (D,F,G) symbols represent individual mice and in (D,G) and bars indicate the mean±SD. Data in (B-E) are representative of two independent experiments and (G) shows combined data from two independent experiments, analyzed by one-way ANOVA followed by Dunnett’s post-test. **: p <0.01.

**Figure S6:**
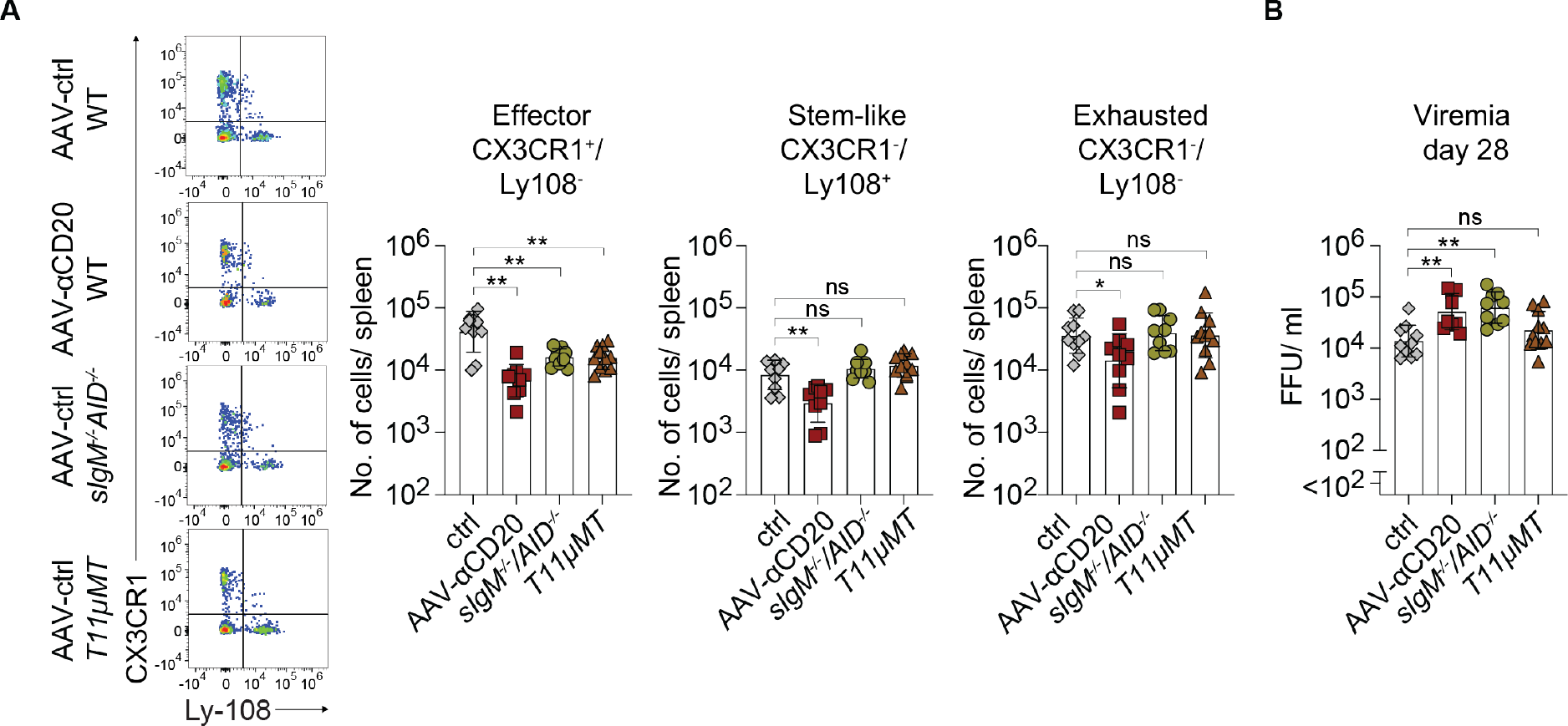
Impact of B cell repertoire-restriction and immunoglobulin-deficiency on virus-specific CD8 T cell subsets. A-B: One week prior to chronic LCMV infection we administered AAV-ctrl to WT, *T11µMT* and to *sIgM^−/−^AID^−/−^*mice and treat WT mice with AAV-αCD20 (same experiment as reported in Fig. 6G-K). Representative FACS plots (A; pre-gated on B220^−^CD4^−^Ter119^−^CD8^+^GP33-Tet^+^ live lymphocytes) and absolute numbers of effector (CX3CR1^+^Ly108^−^), stem-like (CX3CR1^−^Ly108^+^) and exhausted (CX3CR1^−^Ly108^−^) GP33-Tet^+^ CD8 T cells in spleen on d28 after infection. Viremia on day 28 (B). Symbols represent individual mice and bars indicate the mean±SD. Combined data from two independent experiments is shown, analyzed by one-way ANOVA followed by Dunnett’s post-test. *: p<0.05;**: p <0.01; ns: not statistically significant.

